# Modeling the effects of *Ehrlichia Chaffeensis* and movement on dogs

**DOI:** 10.1101/2023.11.28.568825

**Authors:** Folashade B. Agusto, Jaimie Drum

## Abstract

*Ehrlichia chaffeensis* is a tick-borne infectious disease transmitted by *amblyomma americanum* tick. This infectious disease was discovered in the 1970s when military dogs were returning from the Vietnam war. The disease was found to be extremely severe in German Shepards, Doberman Pinschers, Belgium Malinois, and Siberian Huskies. In this study, we developed a mathematical model for dogs and ticks infected with *ehrlichia chaffeensis* with the aim of understanding the impact of movement on dogs as they move from one location to another. This could be a dog taken on a walk in an urban area or on a hike in the mountains. We carried out a global sensitivity analysis with and without movement between three locations using as response functions the sum of acutely and chronically infected and the sum of infected ticks in all life stages. The parameters with the most significant impact on the response functions are dogs disease progression rate, dogs chronic infection progression rate, dogs recovery rate, dogs natural death rate, acutely and chronically infected dogs disease induced death rate, dogs birth rate, eggs maturation rates, tick biting rate, dogs and ticks transmission probabilities, ticks death rate, and the location carrying capacity. Our simulation results show that infection in dogs and ticks are localized in the absence of movement and spreads between locations with highest infection in locations with the highest rate movement. Also, the effect of the control measures which reduces infection trickles to other locations (trickling effect) when control are implemented in a single location. The trickling effect is strongest when control is implemented in a location with the highest movement rate into it.

## 1 Introduction

*Ehrlichia chaffeensis* is a tick-borne infectious disease that lives within white blood cells. The disease was first recognized in Africa during the 1930s. This infectious disease originated in the United States in the 1970s when military dogs were returning from the Vietnam war. It is estimated that 200-250 million dogs died from this disease in Vietnam. Due to the high infection rates of dogs a majority of the war dogs were left behind in Vietnam [20]. They found this disease to be extremely severe in German Shepards, Doberman Pinschers, Belgium Malinois, and Siberian Huskies. This disease can also be referred to as tracker dog disease and tropical canine pancytopenia [21].

One primary vector of *ehrlichia chaffeensis* is the *amblyomma americanum* tick. It is mostly found in south-central and eastern United States [11]. It is also referred to as the Lone Star tick. The tick goes through four different life stages. These life stages are egg, larvae, nymph, and adult, which typically take two years to complete [19]. The adult ticks are typically the ones that feed on dogs. They are seen active March through August in the Midwest [38]. There is currently no vaccine for dogs in preventing *ehrlichia Chaffeensis*. The way of prevention is by preventing ticks on pets and in yards.

There are two stages of infection with *ehrlichia chaffeensis* in dogs. The first 2-4 weeks of infection is referred to as the acute stage. During this stage symptoms include, fever, swollen lymph nodes, respiratory distress, weight loss, bleeding disorders, and sometimes neurological disturbances. There is an antibody and PCR test for *ehrlichia chaffeensis* in dogs. The antibody test could remain negative during the first week of illness and could remain positive for months to years. There is a rapid antibody test that gives results within minutes or there is a ELISA and IFA antibody test that takes days or weeks for results. The PCR test can provides species-specific results and can take days for results. Dogs are treated for *ehrlichia chaffeensis* with a 28-day treatment of doxycycline. After 24-72 hours of initiating treatment the clinical signs should resolve [10]. After 2-4 weeks the dog will recover or progress into the clinical/chronic stage. During the clinical/chronic stage dogs may develop symptoms such as, anemia, bleeding episodes, lameness, eye problems, neurological problems, and swollen limbs. The clinical/chronic stage typically leads to death [21]. There is no correlation between the age of dog and the severity of the infection stage [32]. Dogs are capable of being reinfected of *ehrlichia chaffeensis*, however, dogs cannot pass the disease to their offspring.

To prevent dogs from been infected with *ehrlichia chaffeensis* it is advised to use proper tick prevention techniques. This may include collars, monthly topical, or oral preventatives year round. It is not recommended to use human insect repellent on dogs. After dogs have spent time outside it is recommended to thoroughly evaluate them for ticks. Removing a tick right away reduces the risk of disease transmission. To remove a tick it is recommended to use fine-tipped tweezers and grab the tick closely to the skin’s surface. Then proceed to pull with a steady pressure upwards [10].

The goal of this study is to produce a mathematical model to better understand the transmission of *erhlichia chafeensis* between dogs and *amblyomma americanum*. This model will also be used to better understand the impact of movement on dogs as they move from one location to another. This could be taking a dog on a walk in an urban area cor taking a dog on a hike in the mountains. The rest of the paper is organized as follows: in Section 2, we present the *erhlichia chafeensis* mathematical model for a single location with no movement, the basic qualitative analysis of the model including results of the positivity and boundedness of solutions, the computation of the reproduction number, and global sensitivity analysis. In section 3, we introduce the *erhlichia chafeensis* model with movement between different locations. In this section, we also implement a global sensitivity analysis in the absence and presence of movement. Using results from the sensitivity analysis we simulate the model with movement to determine the effect of disease transmission and ticks natural death when the locations are isolated and when they are connected by movement. In Section 4, we discuss and give conclusions of the results obtained from this study.

## 2 Model formulation

This transmission model of *Ehrlichia Chaffeensis* by incorporating two subgroups: dogs and ticks. The dog population is divided into susceptible (*S*_*D*_), exposed (*E*_*D*_), acute infection (*A*_*D*_), chronic infection (*C*_*D*_), and recovered (*R*_*D*_). Therefore the total population of dogs is given by

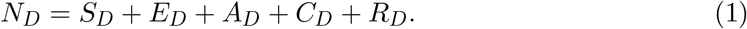

The tick population was divided into egg, larvae, nymph and adult ticks classes. Each of these contained a susceptible class (*S*_*Ti*_) and infections class (*I*_*Ti*_) where *i* = *E, L, N*, and *A*, for egg, larvae, nymph and adult classes. As previously mentioned *ehrlichia chaffeensis* is not transmitted to offspring, so the tick eggs do not have an infection class. Therefore, the total tick population is defined as

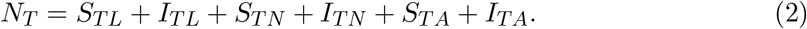

The susceptible dog (*S*_*D*_) compartment is increased due to an increase in newborn dogs. The susceptible (*S*_*D*_) dogs compartment decreases due to a natural death rate (*μ*_*D*_). The force of infection in dogs is represented as

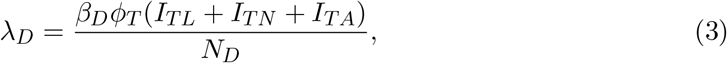

where the parameter *β*_*D*_ is the probability of dog transmission. The parameter *ϕ*_*T*_ represents the rate at which a tick bites a dog. It is assumed that the ticks are all biting at a constant rate. The susceptible dogs progress out of the susceptible compartment at a rate (*λ*_*D*_) to the exposed compartment. The equation for the susceptible dog population compartment is given as

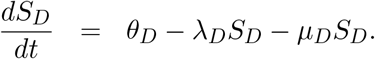

The population of exposed dogs increases at a rates f *λ*_*D*_ and *ελ*_*D*_. These rates comes from infected susceptible dogs and the reinfection of dogs that have recovered from *ehrlichia chaffeensis*. It is found that dogs can only recover from the acute stage of infection since chronic stage illnesses lead to death [23]. Population this compartment decreases due to natural death rate (*μ*_*D*_) and disease progression at the rate (*σ*_*D*_). The exposed dogs compartment is represented by the equation

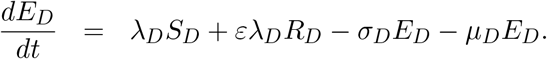

The acute stage compartment (*A*_*D*_) increases from the exposed dog compartment and decreases at a natural death rate of *μ*_*D*_ and an infectious death rate of *δ*_*D*_. It can also decrease at a recovery rate of *γ*_*D*_ or at a progression of infection into the chronic stage at a rate of *υ*_*D*_. The acute stage of the infection in dogs (*A*_*D*_) can be represented as

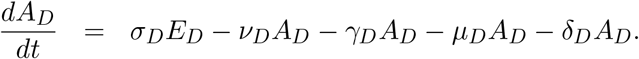

The population of dogs in the chronic stage compartment (*C*_*D*_) increases from the acute stage compartment. This compartment can only decrease due to natural death (*μ*_*D*_) or disease induced death at the rate *δ*_*D*_. The chronic stage of this illness can be managed in dogs, however, it eventually will lead to death, therefore, it does not lead to the recovery of the dogs [23]. The chronic stage (*C*_*D*_) is represented by the following equations

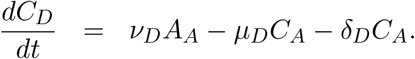

The last compartment of the dog model is the recovered stage (*R*_*D*_). This compartment increases from the acute stage of the disease. At the recovered stage the dog is no longer infectious, therefore, it doesn’t contain an infection death rate. It decreases due to natural death rate (*μ*_*D*_) or due to a reinfection in the dog (*ελ*_*D*_). The recovered stage compartment (*R*_*D*_) is represented as

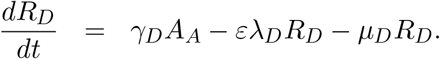

*Amblyomma americanum* ticks transmit *ehrlichia chaffeensis* transstadially (eggs to larvae to nymph to adults) but not transovarially (adults to eggs) [34]. The susceptible ticks move to infectious tick at a rate *λ*_*T*_ which is represented as

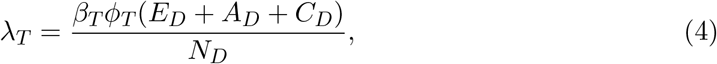

where the parameter *β*_*T*_ represents the probability of tick transmission. The parameter *ϕ*_*T*_ represents the rate at which the ticks bite. The susceptible and infection tick larvae have a maturation rate of *σ*_*L*_ and a natural death rate of *μ*_*L*_. These equations are represented as

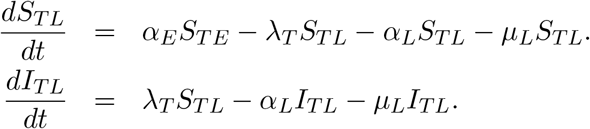

The maturation rate from nymphs to adults is *α*_*N*_ and a natural death rate of *μ*_*N*_. These equations are represented as

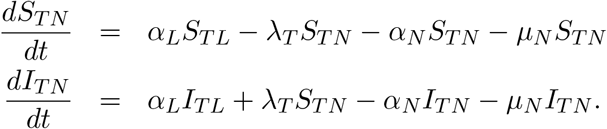

The susceptible and infected adult ticks (*S*_*T A*_ and *I*_*T A*_) lay eggs at a rate of *θ*_*A*_ which is limited by the carrying capacity *K*. Therefore the susceptible tick eggs and susceptible tick adults are represented as

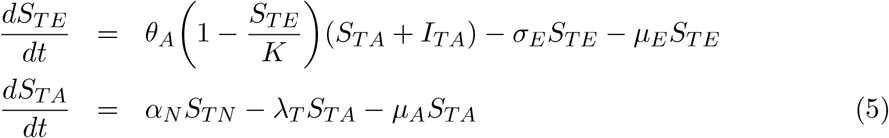

Lastly, infected adults tick (*I*_*T A*_) can only decrease at a death rate of *μ*_*A*_. This is represented as

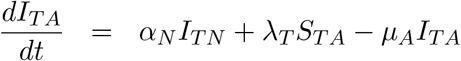

Given the assumptions above, the following nonlinear equations are given for the transmission of *ehrlichia chaffeensis*:

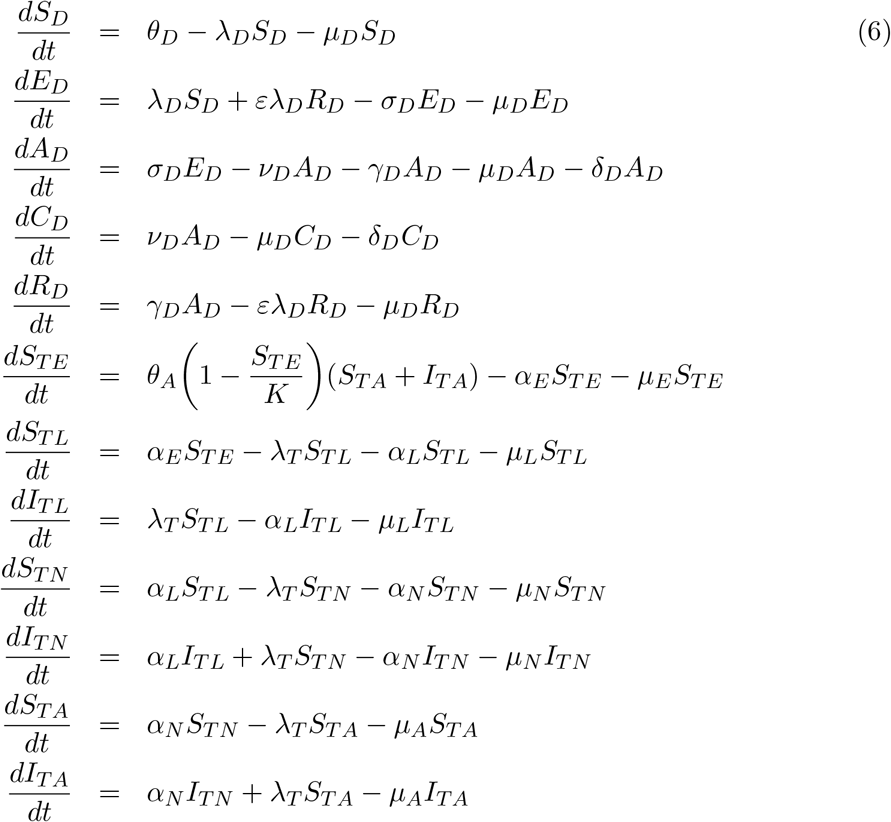

where

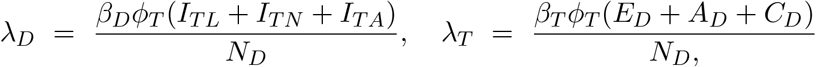

and *N*_*D*_ = *S*_*D*_ + *E*_*D*_ + *A*_*D*_ + *C*_*D*_ + *R*_*D*_.

The conceptualized flow diagram of the *ehrlichia chaffeensis* transmission in dogs model is shown in Figure 1. The corresponding parameters and variables are described in Table 1.

**Table 1:** Description of the parameters and variables for the *erhlichia chaffeensis* model (6).

| Variable | Description |
| --- | --- |
| $S_D$ | Population of susceptible dogs |
| $E_D$ | Population of exposed dogs |
| $A_D$ | Population of acute stage of infection in dogs |
| $C_D$ | Population of chronic stage of infection in dogs |
| $R_D$ | Population of recovered dogs |
| $S_{TE}$ | Susceptible tick eggs |
| $S_{TL}, S_{TN}, S_{TA}$ | Population of susceptible larvae, nymphs, and adult ticks |
| $I_{TE}$ | Infected tick eggs |
| $I_{TL}, I_{TN}, I_{TA}$ | Population of Infected larvae, nymphs, and adult ticks |
| Parameter | Description |
| $\theta_D$ | Birth rate of dogs |
| $\gamma_D$ | Recovery rate in dogs |
| $v_D$ | Acute to chronic disease progression rate in dogs |
| $\sigma_D$ | Disease progression rate in dogs |
| $\varepsilon$ | Reinfection rate of in dogs |
| $\alpha_E$ | Tick eggs to larvae maturation rate |
| $\alpha_L$ | Larvae to nymphs maturation rate |
| $\alpha_N$ | Nymphs to adult ticks maturation rate |
| $\theta_A$ | Ticks egg laying rate |
| $\mu_E$ | Tick eggs decay rate |
| $\mu_L, \mu_N, \mu_A$ | Larvae, nymphs, adult ticks death rate |
| $\mu_D$ | Natural death rate of dogs |
| $\delta_D$ | Acute infection death rate in dogs |
| $\delta_C$ | Chronic infection death rate in dogs |
| $\beta_D$ | Dog transmission probability |
| $\phi_T$ | Tick biting rate |
| $\beta_T$ | Tick transmission probability |

**Figure 1.**
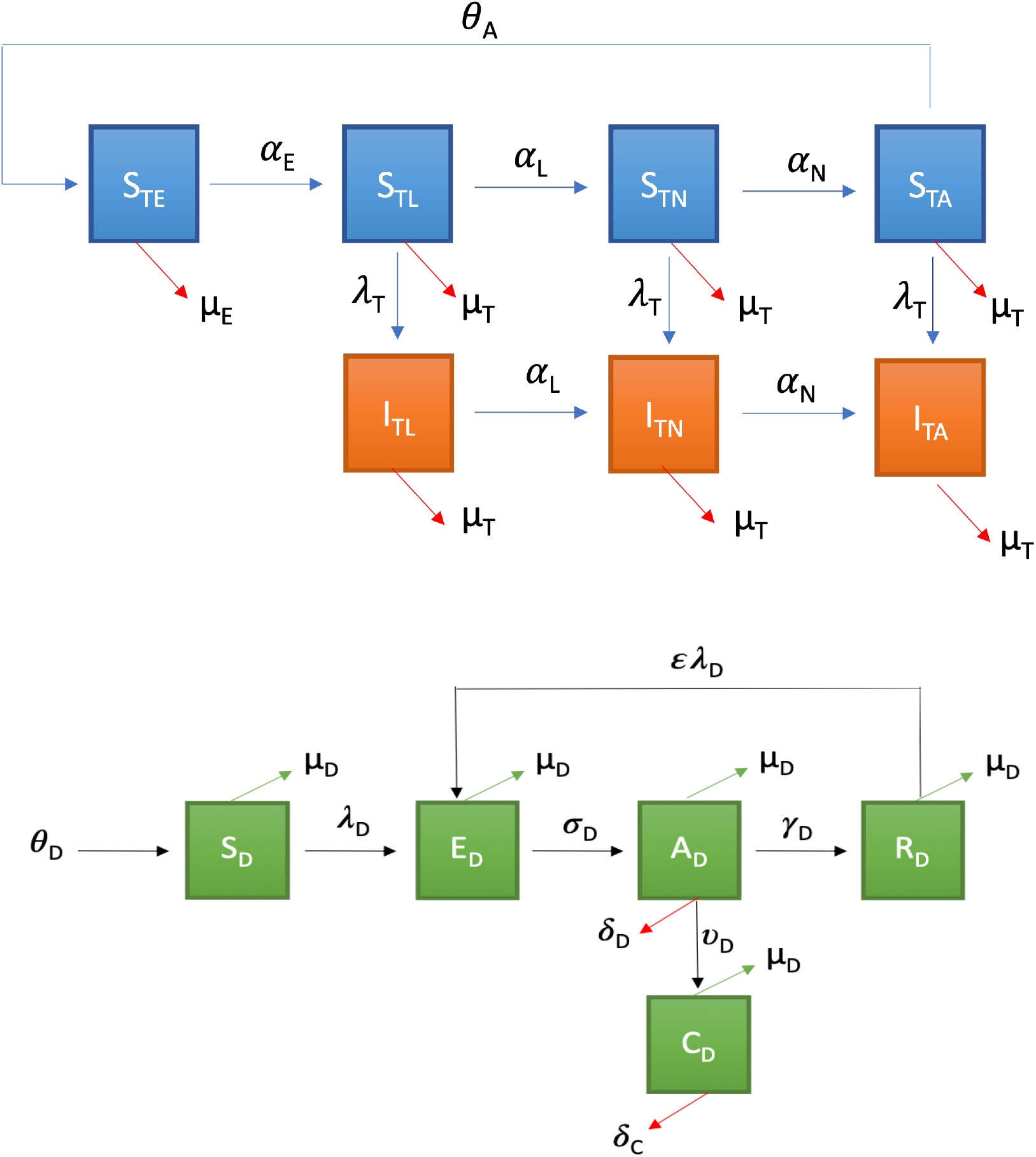
Flow diagram of *erhlichia chaffeensis* model 6. The dogs are divided into susceptible (*S*_*D*_), exposed (*E*_*D*_), acute infection (*A*_*A*_), chronic infection (*C*_*D*_), and recovered (*R*_*D*_). The tick population is composed of susceptible (*S*_*T i*_) and infectious classes (*I*_*T i*_), where *i* = *E, L, N, A* corresponding to egg, larvae, nymph and adult ticks. The blue compartment represent susceptible ticks and the orange compartment represent infected ticks.

### 2.1 Analysis of the model

#### 2.1.1 Basic qualitative properties Positivity and boundedness of solutions

For the Ehrlichia Chaffeensis model (6) to be epidemiologically meaningful, it is important to prove that all its state variables are non-negative for all time. In other words, solutions of the model system (6) with non-negative initial data will remain non-negative for all time *t >* 0.

##### Lemma 1.

*Let the initial data F* (0) *≥* 0, *where F* (*t*) = (*S*_*D*_(*t*), *E*_*D*_(*t*), *A*_*D*_(*t*), *C*_*D*_(*t*), *R*_*D*_(*t*), *S*_*T E*_(*t*), *S*_*T L*_(*t*), *I*_*T L*_(*t*), *S*_*T N*_ (*t*), *I*_*T N*_ (*t*), *S*_*T A*_(*t*), *I*_*T A*_(*t*)). *Then the solutions F* (*t*) *of the Ehrlichia Chaffeensis model* (6) *are non-negative for all t >* 0. *Furthermore*

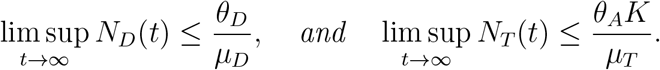

*where*

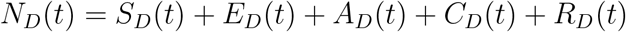

*and*

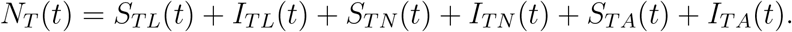

The proof of Lemma 1 is given in Appendix A.

##### Invariant regions

The *ehrlichia chaffeensis* model (6) will be analyzed in a biologically-feasible region as follows. Consider the feasible region

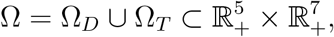

with,

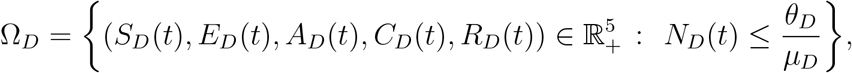

and

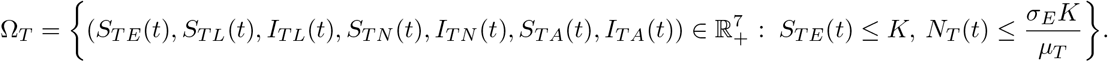

###### Lemma 2.

*The region* 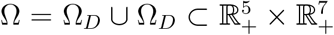 *is positively-invariant for the model* (6) *with non-negative initial conditions in* 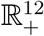.

The prove of Lemma 2 is given in Appendix B.

In the next section, the conditions for the existence and stability of the equilibria of the model (6) are stated.

##### Stability of disease-free equilibrium (DFE)

The *erhlichia chaffeensis* model has a disease-free equilibrium (DFE) denoted by *ε* _0_. The DFE is obtained by setting the right-hand sides of the equations in the model (6) to zero, which is given by

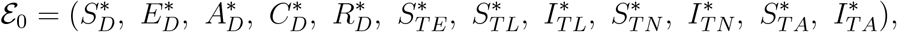

where,

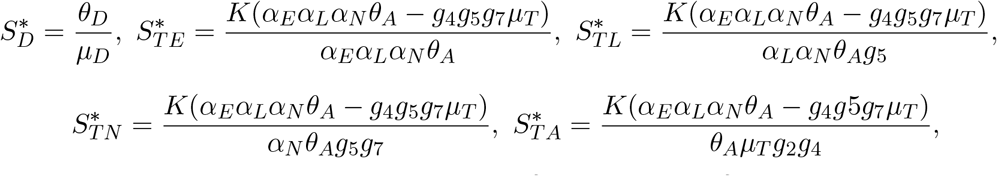

with *g*_1_ = *σ*_*D*_ + *μ*_*D*_, *g*_2_ = *υ*_*D*_ + *γ*_*D*_ + *μ*_*D*_ + *δ*_*D*_, *g*_3_ = *μ*_*D*_ + *δ*_*C*_, *g*_4_ = *σ*_*E*_ + *μ*_*E*_, *g*_5_ = *α*_*L*_ + *μ*_*T*_, *g*_6_ = *α*_*L*_ + *μ*_*T*_, *g*_7_ = *α*_*N*_ + *μ*_*T*_, *g*_8_ = *α*_*N*_ + *μ*_*T*_, and *μ*_*T*_ = min{*μ*_*L*_, *μ*_*N*_, *μ*_*A*_}, and all other disease states are set equal to zero.

The stability of *ε*_0_ can be established using the next generation operator method on system (6). Taking *E*_*D*_, *A*_*D*_, *C*_*D*_, *I*_*T L*_, *I*_*T N*_, and *I*_*T A*_ as the infected compartments and then using the aforementioned notation, the Jacobian *F* and *V* matrices for new infectious terms and the remaining transfer terms, respectively, are defined as:

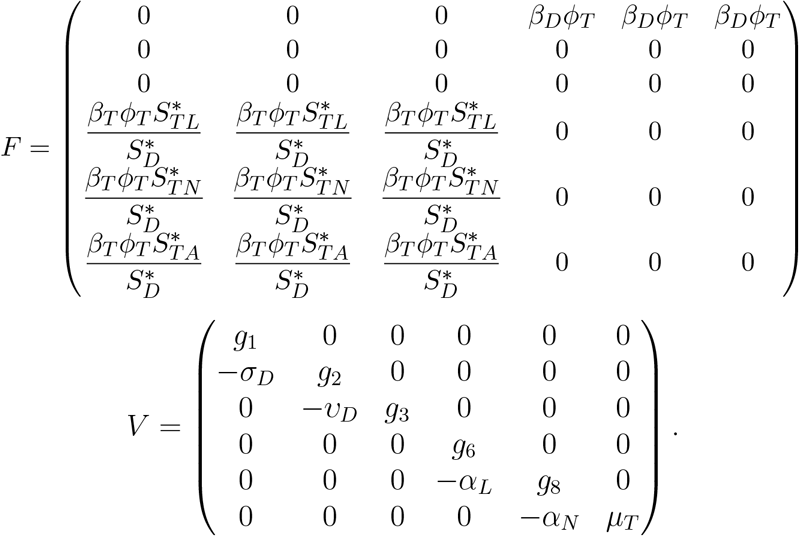

Therefore, using the definition of *ℛ*_0_ = *ρ* (*FV*^− 1^)the *ℛ*_0_ of the model is:

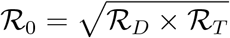

where *ρ* is the spectral radius and

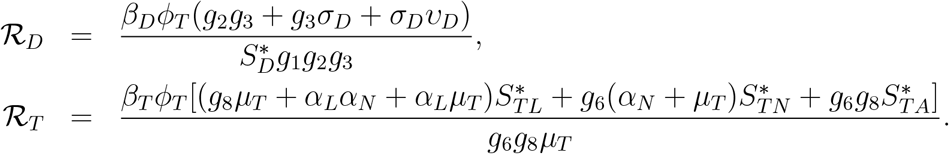

The expression ℛ_*D*_ is the number of secondary infections in dogs. The expressions ℛ_*T*_ is the number of secondary infections in ticks from a single infectious dog. Further, using Theorem 2 in [39], the following result is established.

###### Lemma 3.

*The disease-free equilibrium (DFE) of the ehrlichia chaffeensis model*

(6) *is locally asymptotically stable (LAS) if* ℛ_0_ *<* 1 *and unstable if* ℛ_0_ *>* 1.

The basic reproduction number ℛ_0_ is defined as the average number of new infections that result from one infectious individual (tick or dog) in a population that is fully susceptible. [4, 15, 18, 39]. The epidemiological significance of Lemma 3 is that *ehrlichia chaffeensis* will be eliminated from within a herd if the reproduction number (ℛ_0_) can be brought to (and maintained at) a value less than unity.

### 2.2 Sensitivity analysis of model (6)

In order to determine the contribution of each of the model parameters to key model outputs (such as the number of infected), one can use a sensitivity analysis procedure [8, 13, 16]. Results of the sensitivity analysis help to identify the system parameters that are the best to target during an intervention, and also for future surveillance data gathering. We carried out a global sensitivity analysis using Latin Hypercube Sampling (LHS) and partial rank correlation coefficients (PRCC) to assess the impact of parameter uncertainty and the sensitivity of these key model outputs. The LHS method is a stratified sampling technique without replacement this allows for an efficient analysis of parameter variations across simultaneous uncertainty ranges in each parameter [9, 26, 27, 33]. On the other hand, PRCC measures the strength of the relationship between the model outcome and the parameters, stating the degree of the effect that each parameter has on the model outcome [9, 26, 27, 33].

We start by generating the LHS matrices and assuming all the model parameters are uniformly distributed. We then carry out a total of 1,000 simulations (runs) of the model for the LHS matrix, using the parameter values given in Tables 2 (with ranges varying from *±*20% of the stated baseline values) and as response functions, the sum of carries and infected. The parameter ranking using PRCC are then implemented following these simulation runs.

**Table 2:** Parameter values for each of the parameters for the *ehrlichia chaffeensis* model (6).

| Parameter | Description | Value (1/day) | Reference(s) |
| --- | --- | --- | --- |
| $\theta_D$ | Birth rate of dogs | 70,000 | [29] |
| $\gamma_D$ | Rate of infected to recovered dogs | 0.04761905 | [21] |
| $\nu_D$ | Rate of acute to chronic infection in dogs | 0.04761905 | [21] |
| $\sigma_D$ | Rate of exposed to infected dogs | 0.07142857 | [23] |
| $\varepsilon$ | Rate of reinfection in dogs | 0.00444444 | [25] |
| $\alpha_E$ | Maturation rate from tick eggs to larvae | 0.02439024 | [12] |
| $\alpha_L$ | Maturation rate from tick larvae to nymphs | 0.00273973 | [12] |
| $\alpha_N$ | Maturation rate from nymphs to adult ticks | 0.0037037 | [12] |
| $\theta_A$ | Ticks egg laying rate | 6000 eggs | [6] |
| $\mu_E$ | Death rate of tick eggs | 0.008 | [6] |
| $\mu_T$ | Death rate of ticks | 0.003 | [6] |
| $\mu_D$ | Natural death rate of dogs | 0.00027397 | [5] |
| $\delta_D$ | Acute infection death rate in dogs | 0.1735 | Assumed |
| $\delta_C$ | Chronic infection death rate in dogs | 0.347 | [31] |
| $\beta_D$ | Dog transmission probability | 0.152 | [35] |
| $\phi_T$ | Tick biting rate | 0.044 | Assumed |
| $\beta_T$ | Tick transmission probability | 0.152 | [35] |

The outcome of the global sensitivity analysis is shown in Figure 2 and given in Table 3. The parameters with substantial effect on the sum of the acutely and chronically infected dogs are those parameters whose sensitivity index have significant p-values less than or equal to 0.05. The parameters with the most impacts on (*A*_*D*_ + *C*_*D*_), are disease progression rate in dogs (*σ*_*D*_), the rate (*υ*_*D*_) dogs progress to chronic infection from acute infection, dog recovery rate (*γ*_*D*_), the natural death rate of dogs (*μ*_*D*_), death rate (*δ*_*D*_) of acutely infected dogs, death rate of the chronically infected dogs (*δ*_*C*_), birth rate of dogs (*θ*_*D*_), the maturation rates eggs to larvae (*α*_*E*_), the tick biting rate (*ϕ*_*T*_), the dog transmission probability (*β*_*D*_), the tick transmission probability (*β*_*T*_) and death rate of ticks (*μ*_*T*_), and the carrying capacity *K*. For the sum of infected larvae, nymphs, and adult ticks (*I*_*T L*_ + *I*_*T N*_ + *I*_*T A*_), the significant parameters are *υ*_*D*_, *σ*_*D*_, *μ*_*D*_, *δ*_*D*_, *δ*_*C*_, *μ*_*E*_, *α*_*E*_, *ϕ*_*T*_, *θ*_*D*_, *β*_*T*_, and *μ*_*T*_.

**Table 3:** PRCC and p-values of the *ehrlichia chaffeensis* model (6) using as response functions the sum of infected dogs (*A*_*D*_ + *C*_*D*_) and the sum of infected tick (*I*_*T L*_ + *I*_*T N*_ + *I*_*T A*_) with parameter values in Table 2 that are with *±*20% ranges from the baseline values.

| Parameters | $A_D + C_D$ | | $I_{TL} + I_{TN} + I_{TA}$ | |
| --- | --- | --- | --- | --- |
|  | PRCC | p-value | PRCC | p-value |
| $\varepsilon$ | -0.0335 | 0.2938 | -0.0491 | 0.1242 |
| $\sigma_D$ | -0.3604 | 0.0000 | -0.5323 | 0.0000 |
| $v_D$ | 0.5466 | 0.0000 | 0.2460 | 0.0000 |
| $\gamma_D$ | -0.6476 | 0.0000 | -0.3367 | 0.0000 |
| $\mu_D$ | 0.0711 | 0.0258 | 0.1217 | 0.0001 |
| $\delta_D$ | -0.4754 | 0.0000 | -0.1163 | 0.0003 |
| $\delta_C$ | -0.7851 | 0.0000 | -0.4235 | 0.0000 |
| $\theta_A$ | 0.0051 | 0.8728 | 0.0300 | 0.3478 |
| $\mu_E$ | 0.0184 | 0.5642 | 0.0296 | 0.3544 |
| $\alpha_E$ | 0.7043 | 0.0000 | 0.8301 | 0.0000 |
| $\alpha_L$ | 0.0364 | 0.2548 | 0.0040 | 0.8991 |
| $\alpha_N$ | 0.0117 | 0.7142 | 0.0048 | 0.8815 |
| $\phi_T$ | 0.9484 | 0 | 0.9424 | 0 |
| $\theta_D$ | 0.0893 | 0.0051 | 0.0191 | 0.5488 |
| $\beta_D$ | 0.8800 | 0.0000 | 0.7770 | 0.0000 |
| $\beta_T$ | 0.7142 | 0.0000 | 0.8274 | 0.0000 |
| $\mu_T$ | -0.7243 | 0.0000 | -0.8346 | 0.0000 |
| $K$ | 0.7164 | 0.0000 | 0.8271 | 0.0000 |

**Figure 2.**
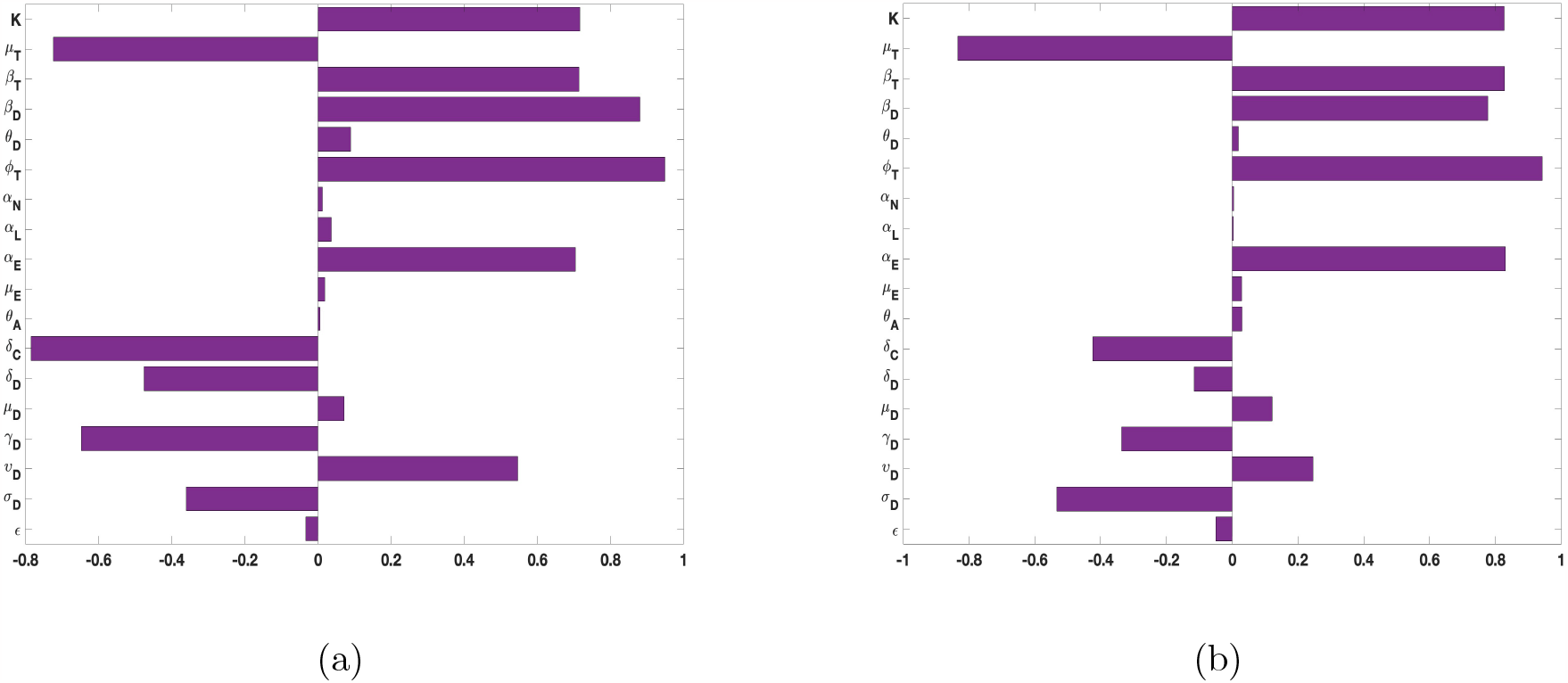
PRCC values for the *ehrlichia chaffeensis* model (6), using as response functions: (a) The sum of infected dogs (*A*_*D*_ + *C*_*D*_); (b) The sum of infected ticks (*I*_*T L*_ + *I*_*T N*_ + *I*_*T A*_). Using parameter values in Table 2 and ranges that *±*20% from the baseline values.

The PRCC index values of some of these parameters have positive signs while others have negative signs. The positive signs means that any increase in these parameters will lead to an increase in the response functions (*A*_*D*_ + *C*_*D*_, and *I*_*T L*_ + *I*_*T N*_ + *I*_*T A*_). While the negative signs implies increase in the parameters will lead to a decrease in the response functions. Hence, these parameters would be useful targets during mitigation efforts.

Therefore, control strategies which targets those parameters with significant PRCC values will give the greatest impact on the model response functions. For instance, a control that aims for a 10% decrease in the transmission probability in dogs (*β*_*D*_) will lead to 88% reduction in infected dogs (*A*_*D*_ + *C*_*D*_) and about 78% reduction in infected ticks (*I*_*T L*_ + *I*_*T N*_ + *I*_*T A*_). Similarly a 10% decrease in the ticks transmission probability (*β*_*T*_) will lead to 71.4% reduced infection in dogs and about 83% reduced infection in ticks. Also, a 10% deduction in tick biting rate (*ϕ*_*T*_) will lead to about 95% reduction in infected dogs, and about 94% reduction in infected ticks. Furthermore, a 10% increase in ticks death rate (*μ*_*T*_) will result in about 72.4% reduction in infected dogs and about 83.4% decrease in ticks.

## 3 Movement model

Next, we extend the *ehrlichia chaffeensis* model (6) by incorporating visitation and long distance migratory movements for dogs and ticks. Dogs are often taken to dog parks or on hiking by their owners, we capture these short dog movement through the visitation parameters *p*_*ij*_, which is the proportion of time a dog in location *i* spends visiting location *j* [1, 37]. Ticks long distance migratory movement may be due to ticks dropping off after feeding on either migratory birds moving north or from white-tail deer or other larger mammals [17]. For the movement of dogs we use Lagrangian model, often use to model short term visitation between places [1, 14, 30]. For ticks movement on the other hand, we model their movement using Eulerian movement [2, 3, 7]. This kind of movement is used to model permanent migratory movement [24, 28, 40]. A number of mathematical models have used this approach to model tick movement [28, 40]. Thus, the movement model is given as

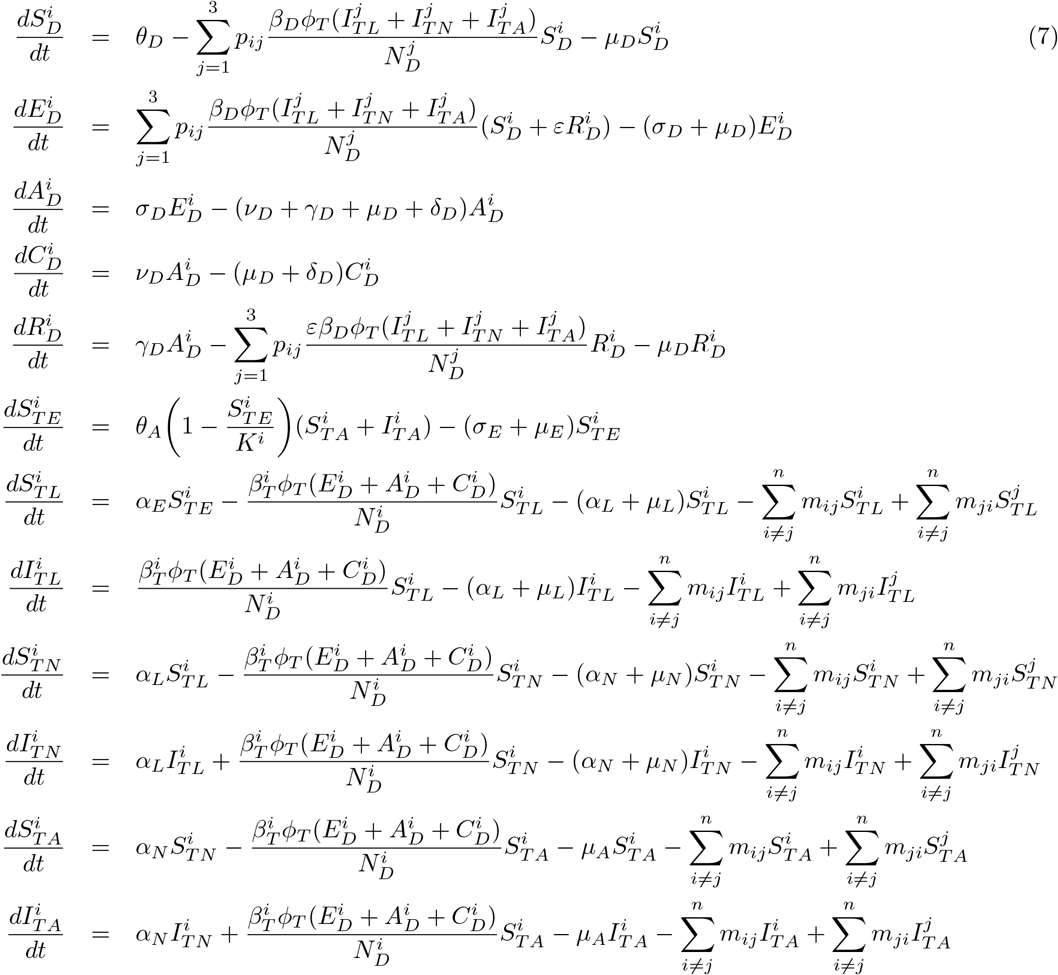

where 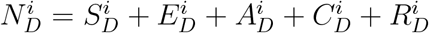. We assumed all the model parameters values are the same in all the locations *i* = 1, · · ·, *n*, except for the carrying capacity(*K*^*i*^), ticks death rate 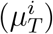, dog and tick transmission probabilities (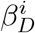and 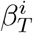). The basic qualitative properties of the *ehrlichia chaffeensis* model (7), the corresponding positivity analysis and the boundedness of solutions are given in Appendix C. The stability analysis of the disease-free equilibrium (DFE) of model (7) leading to the associated reproduction number 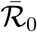 is given in Appendix D.

### 3.1 Sensitivity analysis of model (7) with movement

To carry out global sensitivity analysis for the *ehrlichia chaffeensis* model (7) with movement we also use two response functions, the sum of infected dogs (*A*_*D*_ +*C*_*D*_) and the sum of infected ticks (*I*_*T L*_ +*I*_*T N*_ +*I*_*T A*_). As with the case of the model (6) above, we implemented the sensitivity analysis to determine the contribution of each of the model parameters on these two response functions. We implemented the analysis with three different locations and considered two scenarios, (i) when the locations are isolated; (ii) when the locations are connected with movement of dogs and ticks between them.

To carry out the sensitivity analysis, we fixed some parameters related to the natural history of the infection in dogs and ticks. We then assume the values of parameters like dog birth rate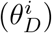, the region carrying capacity (*K*^*i*^), the dogs and ticks transmission probabilities (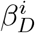, and 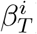), and the ticks death rate 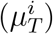 care location dependent since the different locations might have different climatic conditions or micro climates.

Figure 3 (and Table E-1 in Appendix F) show the results of the sensitivity analysis and the p-values of the parameters for the case with no movement between the regions. The results show the parameters with the most impacts o the sum of (*A*_*D*_ + *C*_*D*_)n infected dogs and the sum of infected larvae, nymphs, and adult ticks (*I*_*T L*_ + *I*_*T N*_ + *I*_*T A*_). The significant parameters are disease progression rate in dogs (*σ*_*D*_), the rate (*υ*_*D*_) dogs progress to chronic infection from acute infection, dog recovery rate (*γ*_*D*_), the natural death rate of dogs (*μ*_*D*_), death rate (*δ*_*D*_) of acutely infected dogs, death rate of the chronically infected dogs (*δ*_*C*_), birth rate of dogs 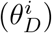, the maturation rates eggs to larvae (*α*_*E*_), the tick biting rate 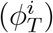, the dog transmission probability (*β*_*D*_), the tick transmission probability 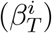 and death rate of ticks 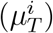, and the carrying capacity *K*^*i*^. The PRCC values of parameters 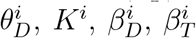, and 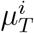are relatively the same since there is no movement between the region.

**Figure 3.**
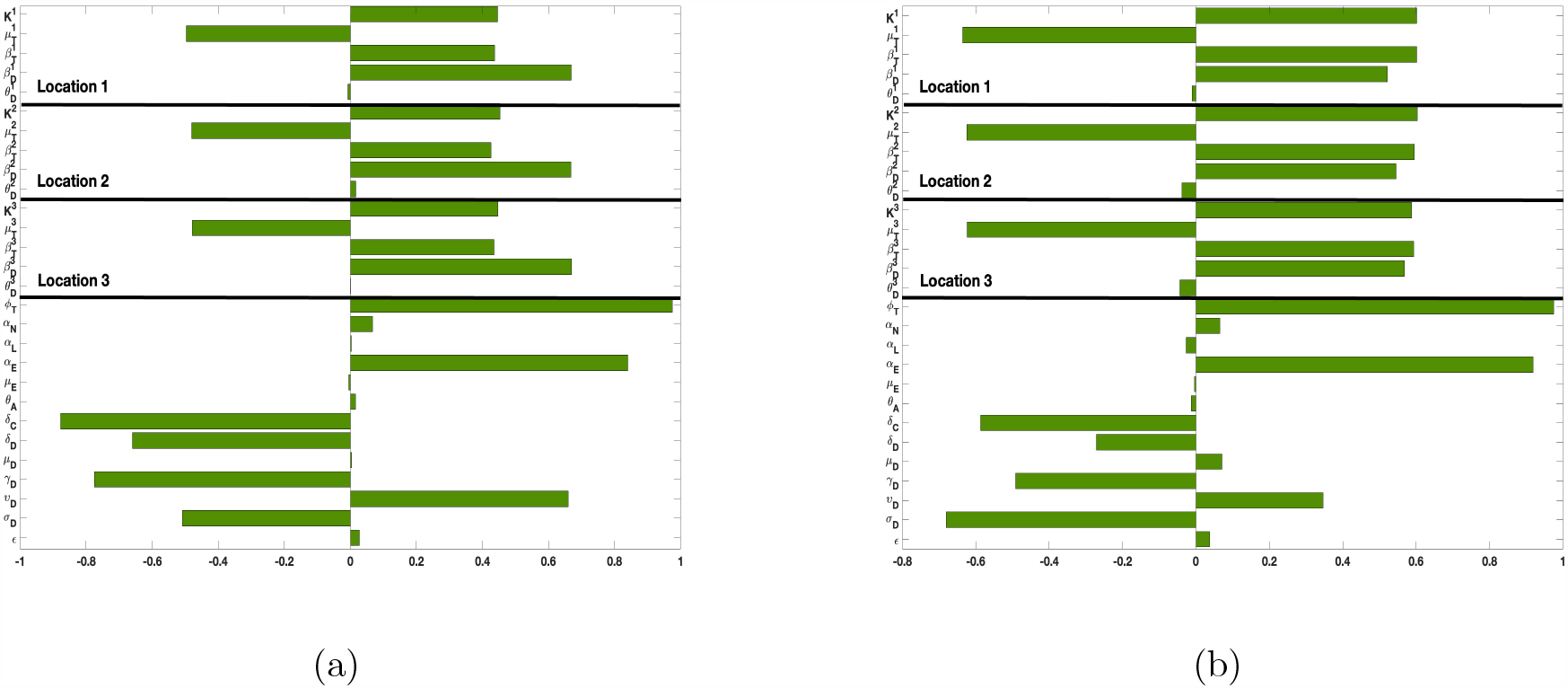
PRCC values for the *ehrlichia chaffeensis* model (7) with no movement between the regions using as response functions: (a) The sum of infected dogs (*A*_*D*_ + *C*_*D*_); (b) The sum of infected ticks (*I*_*T L*_ + *I*_*T N*_ + *I*_*T A*_). Using parameter values in Table 2 and ranges that *±*20% from the baseline values.

Figure 4 (and Table E-2 in Appendix F) show the results of the sensitivity analysis and the p-values of the parameters for the case with movement. The significant parameters *σ*_*D*_, *υ*_*D*_, *γ*_*D*_, *μ*_*D*_, *δ*_*D*_, *δ*_*C*_, *α*_*E*_ are relatively the same since we used the same parameters between the regions. On the other hand, the PRCC values of parameters 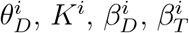, and 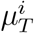 are different, with those in location 1 having higer values than those in location 2 and 3 since there is more movement into location 1, than 2 and 3.

**Figure 4.**
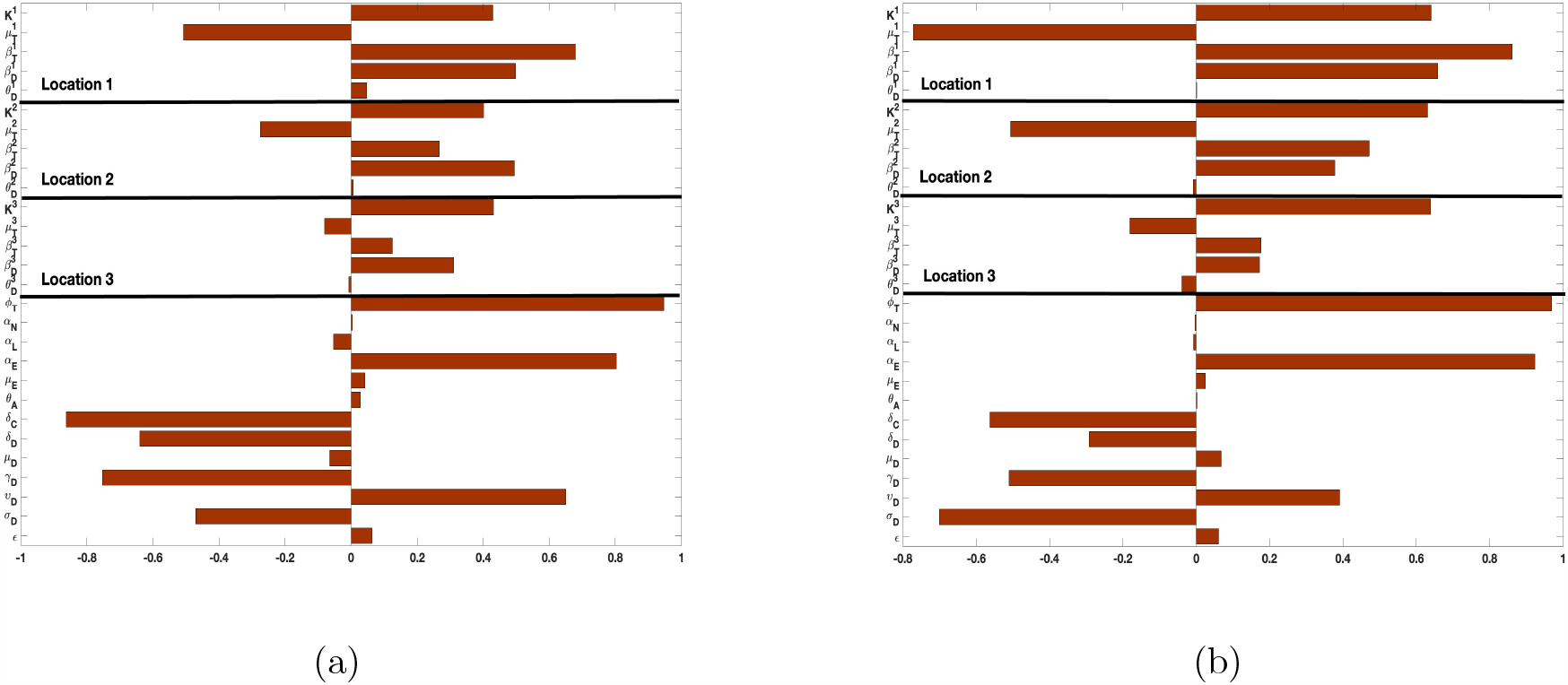
PRCC values for the *ehrlichia chaffeensis* model (7) with movement between the regions using as response functions: (a) The sum of infected dogs (*A*_*D*_ + *C*_*D*_); (b) The sum of infected ticks (*I*_*T L*_ + *I*_*T N*_ + *I*_*T A*_). Using parameter values in Table 2 and ranges that *±*20% from the baseline values. Under this scenario, there is increased movement to location 1, than 2 and 3. Location 3 has the least movement into it.

### Simulating the *ehrlichia chaffeensis* model (7)

In this section, we would simulate the *ehrlichia chaffeensis* model (7) when there is no movements between the three locations and when dogs and ticks move between the locations. Later on, we would use the results from the sensitivity analysis and simulate the *ehrlichia chaffeensis* model (7) varying the transmission probabilities 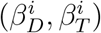, and the ticks death rate 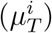. Then we would analyze the effect of these parameters on the spread of *ehrlichia chaffeensis* separately and jointly.

We start by simulating model (7) with no movement using parameters given in Table 2. We assume that infection is higher in location 1 followed by location 2, and location 3 has the least infection. As expected Figures 5(a) and 5(b) shows higher number of acutely and chronically infected dogs in location 1 followed by locations 2 and 3; higher infected larvae, nymphs adult ticks were also observed in Figures 5(c), 5(d), and 5(e) in location 1 followed by locations 2 and 3.

**Figure 5.**
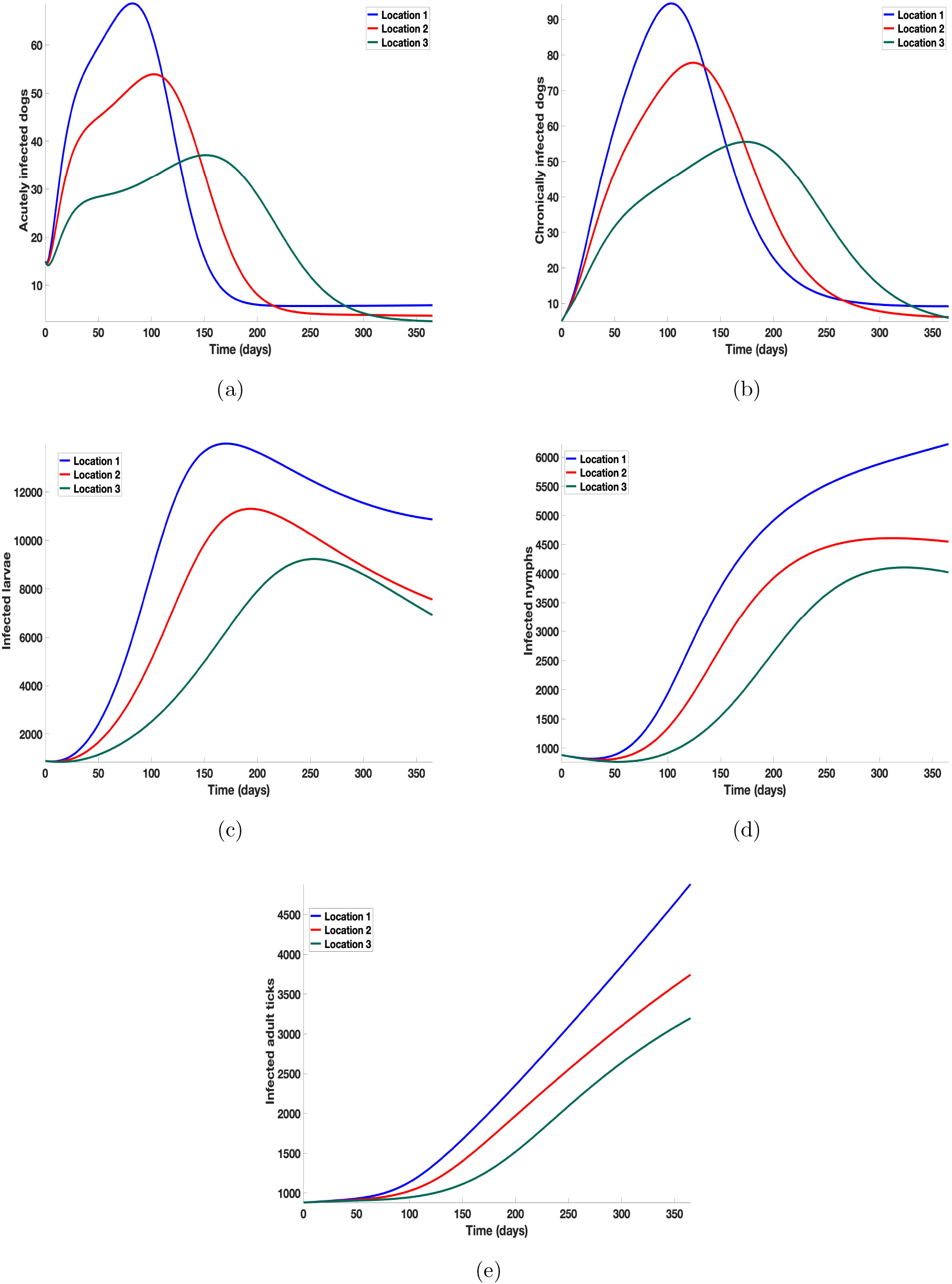
Simulation results of model (7) with no movement using parameters given in Table 2. Infection is higher in location 1 followed by location 2, location 3 has the least infection. (a) acutely infected dogs; (b) chronically infected dogs; (c) infected larvae (d) Infected nymphs (e) infected adult ticks.

Next, we simulate model (7) with more movement into location 3 and least movement into locations 1 while infection remain higher in location 1 and smallest in location 3. With this scenario, Figures 6(a) and 6(b) shows higher number of acutely and chronically infected dogs in location 3 followed by locations 2 then 1 even though infection is higher in location 1. Also, higher infected larvae, nymphs adult ticks were observed in location 3 followed by locations 2 and 1, see Figures 6(c), 6(d), and 6(e).

**Figure 6.**
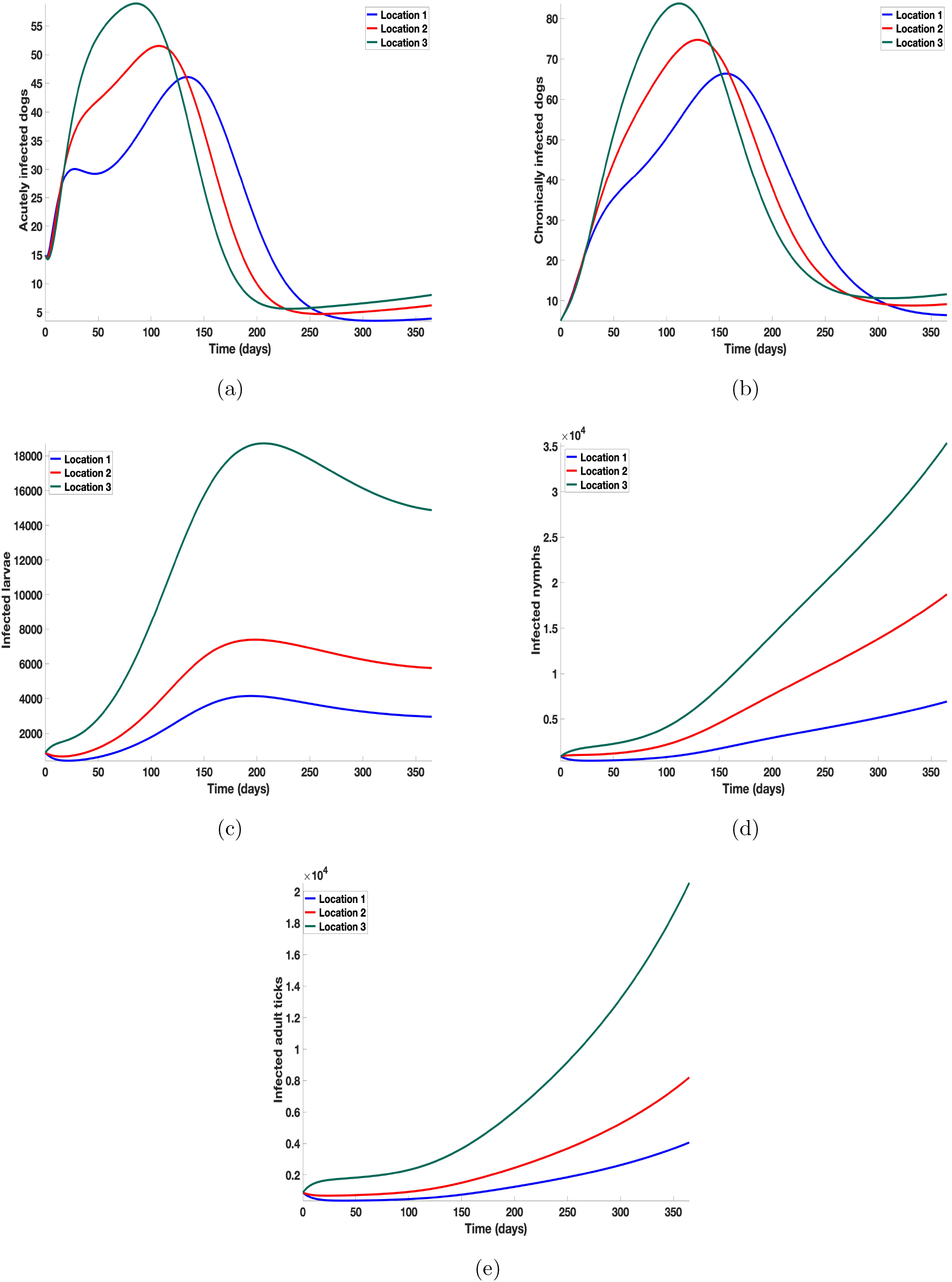
Simulation results of model (7) with movement using parameters given in Table 2. Infection is higher in location 1 followed by location 2, location 3 has the least infection. More movement to location 3 from locations 1, and 3. (a) acutely infected dogs; (b) chronically infected dogs; (c) infected larvae (d) infected nymphs (e) Infected adult ticks.

These results shows the impact of movement on the transmission of the disease as dogs and ticks move across regions.

In the next section we would use the results from the sensitivity analysis and simulate the *ehrlichia chaffeensis* model (7) varying the transmission probabilities 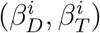 and the ticks’ death rate 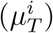. We analyze jointly the effect of these parameters on the spread of *ehrlichia chaffeensis* within the dogs and ticks populations.

### Effect of disease transmission and ticks natural death

Here we investigate the impact of disease transmission probabilities and the ticks death rate as control measures on infected dogs and ticks. We vary the values of the transmission probabilities 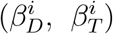and the number of ticks death rate 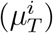 and then examine the effect of these measures separately and jointly on the trajectories of infected dogs and ticks under several scenarios (i) when the three locations are isolated with no movement between them and control measures are implemented in each location; (ii) when the locations are connected with movement between and control measures are first implemented a location at a time and then implemented at once in all the locations.

### Isolated locations: Disease transmission and ticks control

Here we explore the combined effect of varying the transmission probabilities (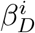, and 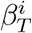) and ticks natural death rate 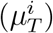 when there are no movements between the locations. However, infection is higher in location 1 followed by location 2, location 3 has the least infection. We considered three measures, (i) measure 1 where the baseline parameters are used for 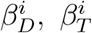, and 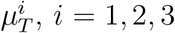, as given in Table 2; (ii) measure 2, the baseline parameter values for 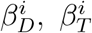, are halved while the value for 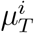 is doubled; (iii) measure 3, the baseline parameter values for 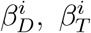 are divided by four while the values for 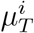are multiplied by four.

In Figure 7, we observed reduction in acutely and chronically infected dogs in each location as the control measures varies as described above; Figure F-1 in Appendix G show the infected larvae, nymphs, and adult ticks in each locations. Table 4 show similar trends with the sum of the acutely and chronically infected dogs and ticks over the simulation period of 52 weeks representing a year.

**Table 4:**
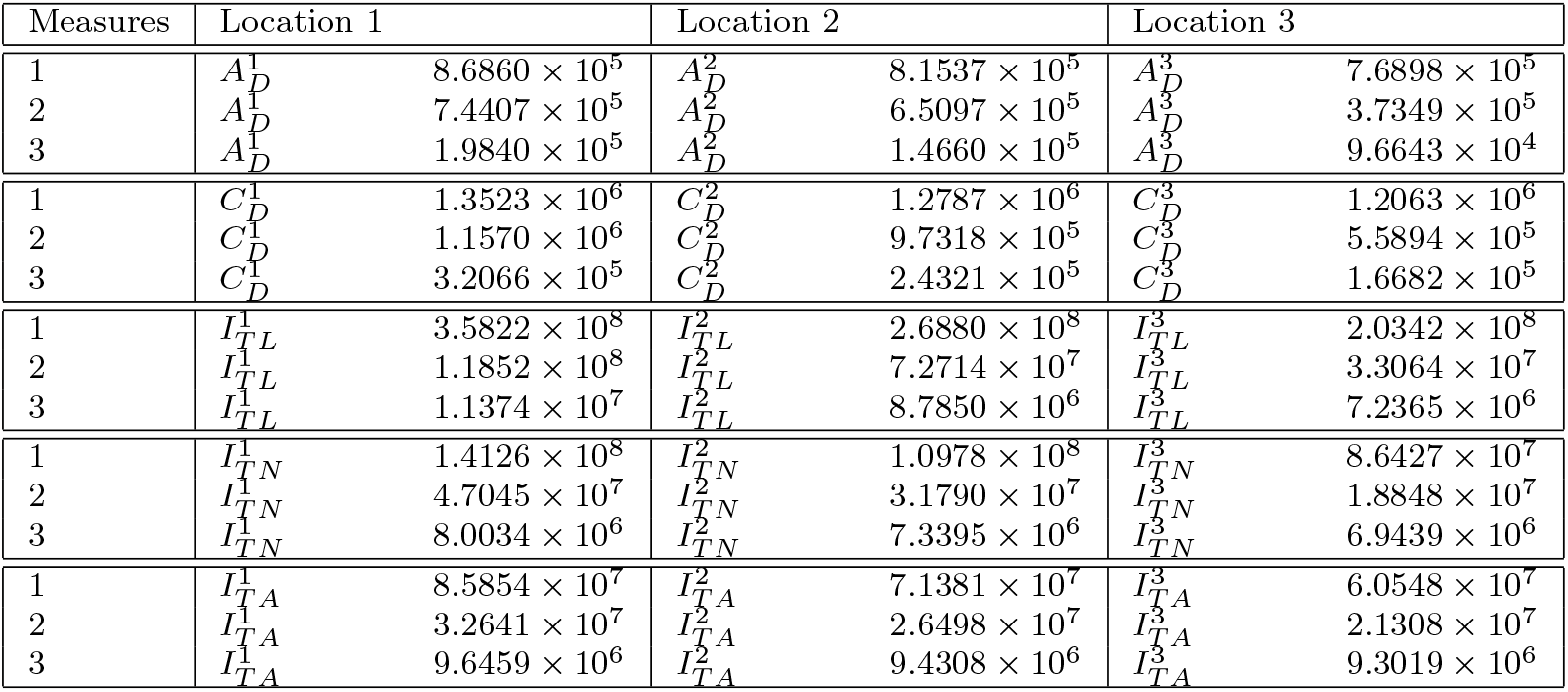
Sum of the simulation of model (7) with no movement into the different locations using three different control measures. Measure 1: used the baseline parameters for 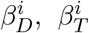, and 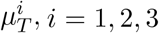,, as given in Table 2; Measure 2: the parameters for 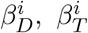, are divided by two and 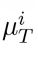 are doubled; Measure 3: 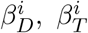, values are divided by four and 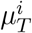 are multiplied by four.

**Figure 7.**
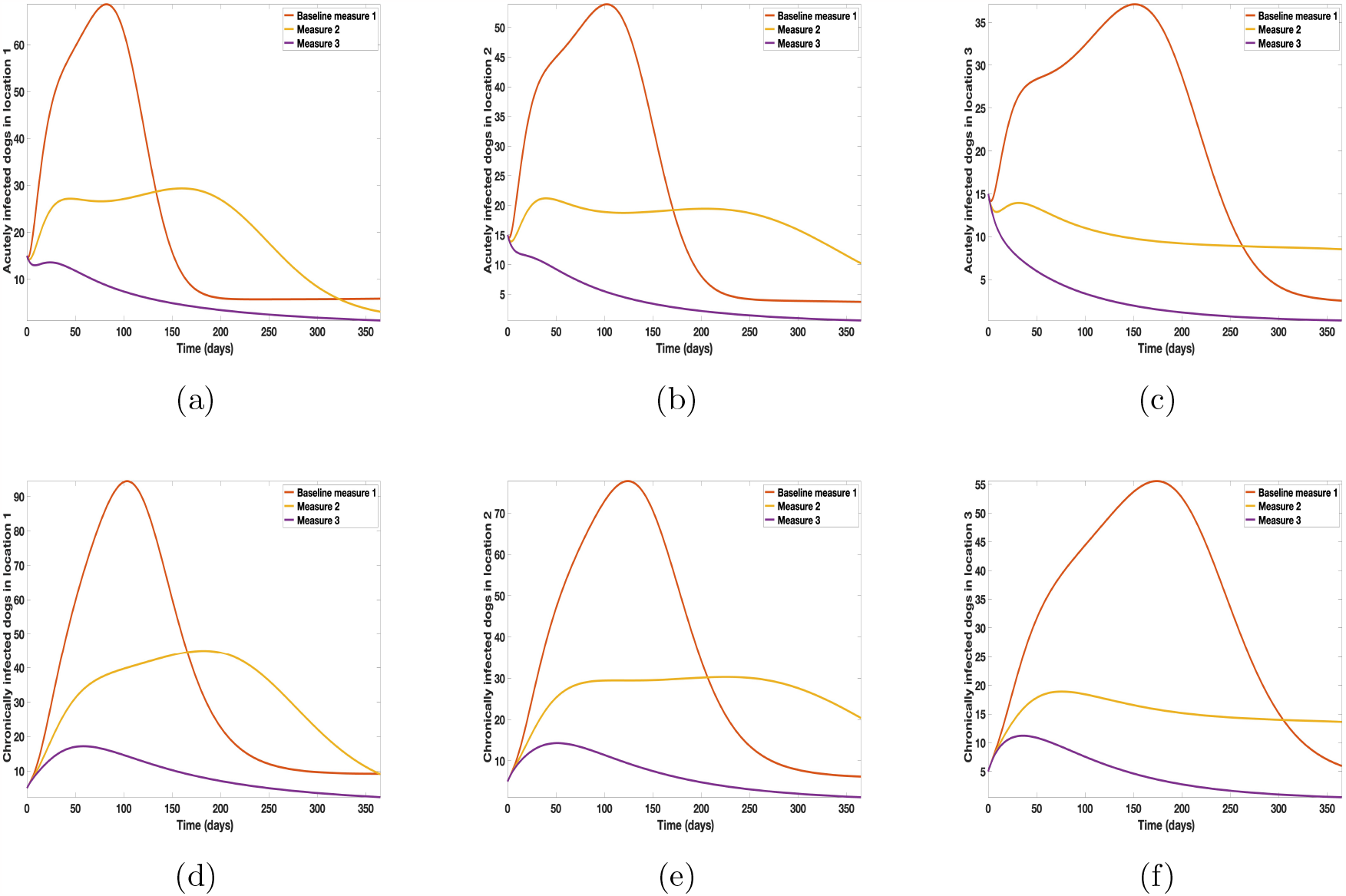
Simulation results of model (7) with no movement using parameters given in Table 2. Infection is higher in location 1 followed by location 2, location 3 has the least infection. More movement to location 3 from locations 1, and 3. (a) - (c) acutely infected dogs; (d) - (f) chronically infected dogs.

This results show the importance of ensuring the infection rates are low in order to reduce the overall burden of the disease in each location.

### Connected locations: Control in one location at a time

In this section, we explore the effect of varying the transmission probabilities (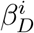, and 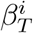) and ticks natural death rate 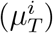 when dogs and ticks can move freely between the locations. We assume the infection is higher in location 1 followed by locations 2, and 3; location 3 has the least infection. Here we consider the scenario where the control measures are implemented a location at a time. We also considered three control measures 1,2, 3: with measure 1, the baseline parameter values are used for 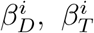, and 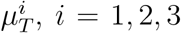, as given in Table 2; with measure 2, the baseline parameter values for 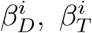, are halved while and 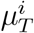 is doubled; with measure 3, the baseline parameter values for 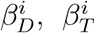 are divided by four while 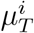are multiplied by four.

### Connected locations: Control in location 1 only

With this scenario the control measures described above are only implemented location 1. We observed in location 1, substantial reduction in population of acutely and chronically infected dogs as the control measures varies from Measures 1 through 3, see Figure 8. The effect of this measure trickles to locations 2 and 3 but the impact is minimal compare to the effect in location 1. Figure F-2 in Appendix G.1 shows the outcome for infected larvae, nymphs, and adult ticks in each locations. Table 5 shows similar trends in the sum of the acutely and chronically infected dogs and ticks over the simulation period.

**Table 5:** Sum of the simulation results of model (7) with movement. Control measures are only implemented in location 1.

| Measures | Location 1 |  | Location 2 |  | Location 3 |  |
| --- | --- | --- | --- | --- | --- | --- |
| 1 | $A_D^1$ | $7.8998 \times 10^5$ | $A_D^2$ | $8.2910 \times 10^5$ | $A_D^3$ | $8.6320 \times 10^5$ |
| 2 | $A_D^1$ | $7.3806 \times 10^5$ | $A_D^2$ | $8.1344 \times 10^5$ | $A_D^3$ | $8.4190 \times 10^5$ |
| 3 | $A_D^1$ | $5.7417 \times 10^5$ | $A_D^2$ | $7.9930 \times 10^5$ | $A_D^3$ | $8.2301 \times 10^5$ |
| 1 | $C_D^1$ | $1.2379 \times 10^6$ | $C_D^2$ | $1.2905 \times 10^6$ | $C_D^3$ | $1.3359 \times 10^6$ |
| 2 | $C_D^1$ | $1.1365 \times 10^6$ | $C_D^2$ | $1.2705 \times 10^6$ | $C_D^3$ | $1.3089 \times 10^6$ |
| 3 | $C_D^1$ | $8.3462 \times 10^5$ | $C_D^2$ | $1.2531 \times 10^6$ | $C_D^3$ | $1.2859 \times 10^6$ |
| 1 | $I_{TL}^1$ | $9.8522 \times 10^7$ | $I_{TL}^2$ | $1.8229 \times 10^8$ | $I_{TL}^3$ | $4.6149 \times 10^8$ |
| 2 | $I_{TL}^1$ | $7.5886 \times 10^7$ | $I_{TL}^2$ | $1.5423 \times 10^7$ | $I_{TL}^3$ | $3.8438 \times 10^8$ |
| 3 | $I_{TL}^1$ | $5.7685 \times 10^7$ | $I_{TL}^2$ | $1.3168 \times 10^6$ | $I_{TL}^3$ | $3.2258 \times 10^8$ |
| 1 | $I_{TN}^1$ | $1.0240 \times 10^8$ | $I_{TN}^2$ | $2.7218 \times 10^8$ | $I_{TN}^3$ | $5.1081 \times 10^8$ |
| 2 | $I_{TN}^1$ | $7.9835 \times 10^7$ | $I_{TN}^2$ | $2.1841 \times 10^8$ | $I_{TN}^3$ | $4.1087 \times 10^8$ |
| 3 | $I_{TN}^1$ | $5.8344 \times 10^7$ | $I_{TN}^2$ | $1.6843 \times 10^8$ | $I_{TN}^3$ | $3.1871 \times 10^8$ |
| 1 | $I_{TA}^1$ | $5.2300 \times 10^7$ | $I_{TA}^2$ | $1.0372 \times 10^8$ | $I_{TA}^3$ | $2.5791 \times 10^8$ |
| 2 | $I_{TA}^1$ | $4.2156 \times 10^7$ | $I_{TA}^2$ | $8.6180 \times 10^7$ | $I_{TA}^3$ | $2.1335 \times 10^8$ |
| 3 | $I_{TA}^1$ | $3.1419 \times 10^7$ | $I_{TA}^2$ | $6.8570 \times 10^7$ | $I_{TA}^3$ | $1.6840 \times 10^8$ |

**Figure 8.**
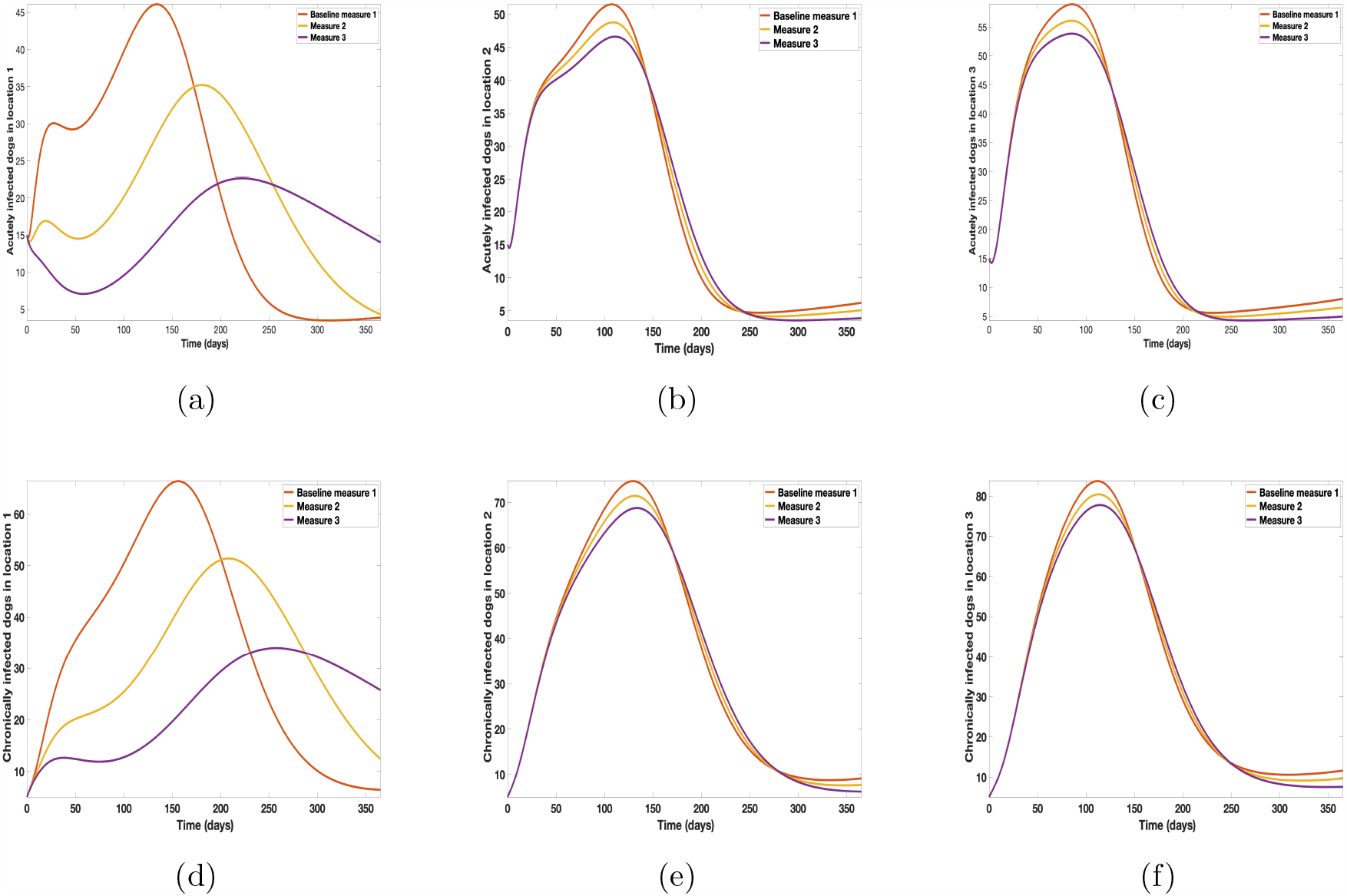
Simulation results of model (7) with movement. Control measures are only implemented in location 1. (a) acutely infected dogs; (b) chronically infected dogs; (c) infected larvae (d) infected nymphs (e) infected adult ticks.

### Connected locations: Control in location 2 only

With this scenario the control measures described above are only implemented in location 2.Significant reduction in population of acutely and chronically infected dogs is observed in location 2 as the control measures varies from Measures 1 through 3, see Figure 9. The effect of this measure trickles to locations 1 and 3 but the impact is minimal compare to the effect in location 2. Figure F-3 in Appendix G.2.1 shows the outcome for infected larvae, nymphs, and adult ticks in each locations. Similar trends can be seen in Table 6 in the values of the sum of the acutely and chronically infected dogs and ticks over the simulation period.

**Table 6:** Sum of the simulation results of model (7) with movement. Control measures are only implemented in location 2.

| Measures | Location 1 |  | Location 2 |  | Location 3 |  |
| --- | --- | --- | --- | --- | --- | --- |
| 1 | $A_D^1$ | $7.8998 \times 10^5$ | $A_D^2$ | $8.2910 \times 10^5$ | $A_D^3$ | $8.6320 \times 10^5$ |
| 2 | $A_D^1$ | $7.7723 \times 10^5$ | $A_D^2$ | $7.6326 \times 10^5$ | $A_D^3$ | $8.3576 \times 10^5$ |
| 3 | $A_D^1$ | $7.6435 \times 10^5$ | $A_D^2$ | $6.5388 \times 10^5$ | $A_D^3$ | $8.1033 \times 10^5$ |
| 1 | $C_D^1$ | $1.2379 \times 10^6$ | $C_D^2$ | $1.2905 \times 10^6$ | $C_D^3$ | $1.3359 \times 10^6$ |
| 2 | $C_D^1$ | $1.2197 \times 10^6$ | $C_D^2$ | $1.1930 \times 10^6$ | $C_D^3$ | $1.3011 \times 10^6$ |
| 3 | $C_D^1$ | $1.1996 \times 10^6$ | $C_D^2$ | $9.6952 \times 10^5$ | $C_D^3$ | $1.2695 \times 10^6$ |
| 1 | $I_{TL}^1$ | $9.8522 \times 10^7$ | $I_{TL}^2$ | $1.8229 \times 10^8$ | $I_{TL}^3$ | $4.6149 \times 10^8$ |
| 2 | $I_{TL}^1$ | $7.9056 \times 10^7$ | $I_{TL}^2$ | $1.3233 \times 10^8$ | $I_{TL}^3$ | $3.6249 \times 10^8$ |
| 3 | $I_{TL}^1$ | $6.2368 \times 10^7$ | $I_{TL}^2$ | $9.1858 \times 10^7$ | $I_{TL}^3$ | $2.7911 \times 10^8$ |
| 1 | $I_{TN}^1$ | $1.0240 \times 10^8$ | $I_{TN}^2$ | $2.7218 \times 10^8$ | $I_{TN}^3$ | $5.1081 \times 10^8$ |
| 2 | $I_{TN}^1$ | $7.7780 \times 10^7$ | $I_{TN}^2$ | $1.9816 \times 10^8$ | $I_{TN}^3$ | $3.8444 \times 10^8$ |
| 3 | $I_{TN}^1$ | $5.4568 \times 10^7$ | $I_{TN}^2$ | $1.2939 \times 10^8$ | $I_{TN}^3$ | $2.6622 \times 10^8$ |
| 1 | $I_{TA}^1$ | $5.2300 \times 10^7$ | $I_{TA}^2$ | $1.0372 \times 10^8$ | $I_{TA}^3$ | $2.5791 \times 10^8$ |
| 2 | $I_{TA}^1$ | $4.0332 \times 10^7$ | $I_{TA}^2$ | $7.6112 \times 10^7$ | $I_{TA}^3$ | $1.9685 \times 10^8$ |
| 3 | $I_{TA}^1$ | $2.8382 \times 10^7$ | $I_{TA}^2$ | $4.9422 \times 10^7$ | $I_{TA}^3$ | $1.3647 \times 10^8$ |

**Figure 9.**
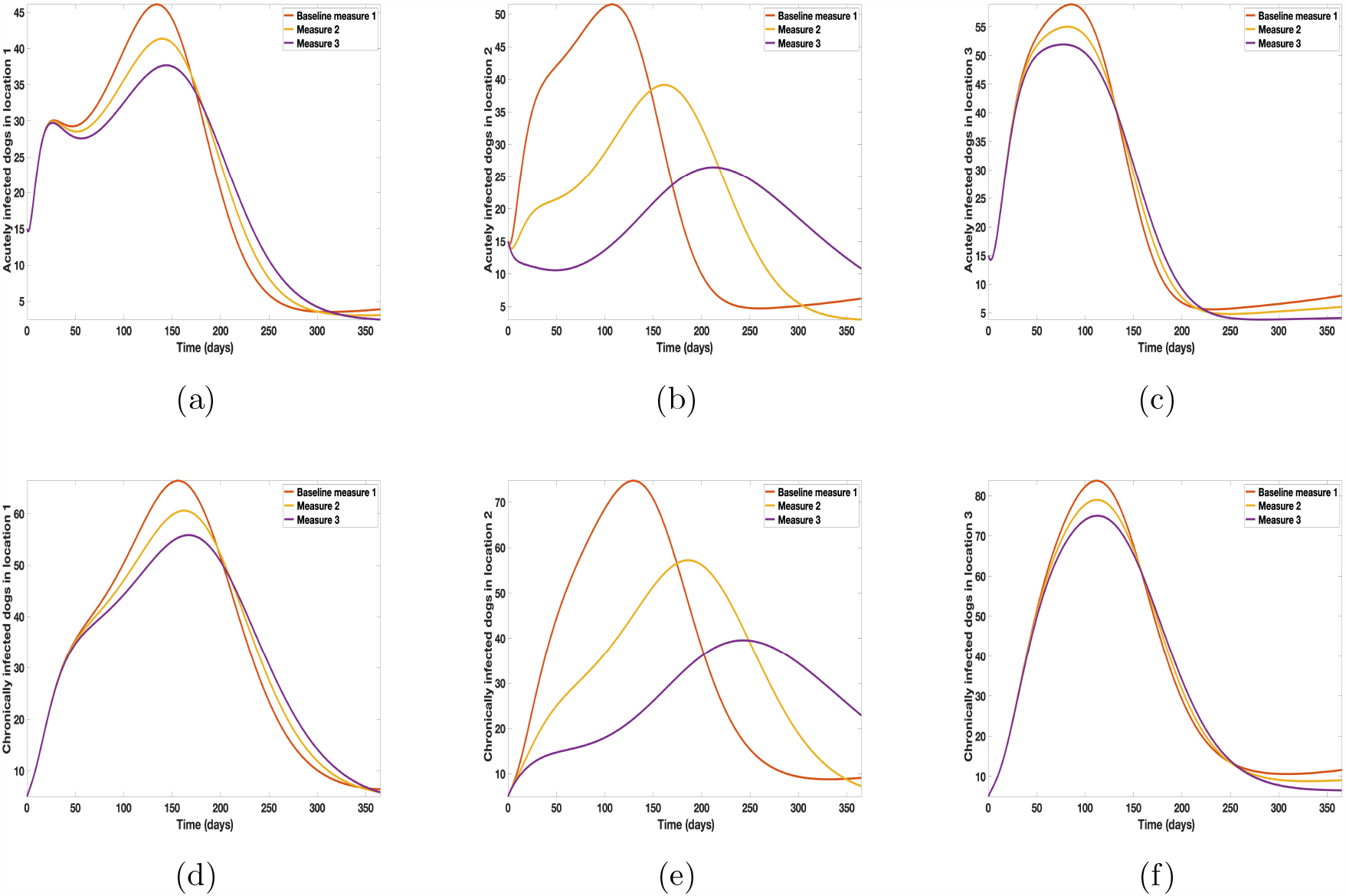
Simulation results of model (7) with movement. Control measures are only implemented in location 2. (a) acutely infected dogs; (b) chronically infected dogs; (c) infected larvae (d) infected nymphs (e) infected adult ticks.

### Connected locations: Control in location 3 only

With this scenario the control measures are implemented in location 3 only. We observed in location 3, significant reduction in population of acutely and chronically infected dogs with changes in the control measures 1 through 3, see Figure 10. The effect of this measure trickles to locations 1 and 2 but the effect is minimal compare to the effect in location 3. Table 7 shows similar trends in the the sum of the acutely and chronically infected dogs and ticks over the simulation period of 52 weeks. The outcome for infected larvae, nymphs, and adult ticks in each locations are shown in Figure F-4 in Appendix G.2.3.

**Table 7:** Sum of the simulation results of model (7) with movement. Control measures are only implemented in location 3.

| Measures | Location 1 |  | Location 2 |  | Location 3 |  |
| --- | --- | --- | --- | --- | --- | --- |
| 1 | $A_D^1$ | $7.8998 \times 10^5$ | $A_D^2$ | $8.2910 \times 10^5$ | $A_D^3$ | $8.6320 \times 10^5$ |
| 2 | $A_D^1$ | $7.6634 \times 10^5$ | $A_D^2$ | $7.9359 \times 10^5$ | $A_D^3$ | $7.6814 \times 10^5$ |
| 3 | $A_D^1$ | $7.2907 \times 10^5$ | $A_D^2$ | $7.6287 \times 10^5$ | $A_D^3$ | $5.8953 \times 10^5$ |
| 1 | $C_D^1$ | $1.2379 \times 10^6$ | $C_D^2$ | $1.2905 \times 10^6$ | $C_D^3$ | $1.3359 \times 10^6$ |
| 2 | $C_D^1$ | $1.2013 \times 10^6$ | $C_D^2$ | $1.2441 \times 10^6$ | $C_D^3$ | $1.2033 \times 10^6$ |
| 3 | $C_D^1$ | $1.1267 \times 10^6$ | $C_D^2$ | $1.1978 \times 10^6$ | $C_D^3$ | $8.6925 \times 10^5$ |
| 1 | $I_{TL}^1$ | $9.8522 \times 10^7$ | $I_{TL}^2$ | $1.8229 \times 10^8$ | $I_{TL}^3$ | $4.6149 \times 10^8$ |
| 2 | $I_{TL}^1$ | $6.2381 \times 10^7$ | $I_{TL}^2$ | $1.1297 \times 10^8$ | $I_{TL}^3$ | $2.6367 \times 10^8$ |
| 3 | $I_{TL}^1$ | $3.5622 \times 10^7$ | $I_{TL}^2$ | $6.3083 \times 10^7$ | $I_{TL}^3$ | $1.2774 \times 10^8$ |
| 1 | $I_{TN}^1$ | $1.0240 \times 10^8$ | $I_{TN}^2$ | $2.7218 \times 10^8$ | $I_{TN}^3$ | $5.1081 \times 10^8$ |
| 2 | $I_{TN}^1$ | $6.1195 \times 10^7$ | $I_{TN}^2$ | $1.6251 \times 10^8$ | $I_{TN}^3$ | $2.9118 \times 10^8$ |
| 3 | $I_{TN}^1$ | $2.9903 \times 10^7$ | $I_{TN}^2$ | $7.9561 \times 10^7$ | $I_{TN}^3$ | $1.3070 \times 10^8$ |
| 1 | $I_{TA}^1$ | $5.2300 \times 10^7$ | $I_{TA}^2$ | $1.0372 \times 10^8$ | $I_{TA}^3$ | $2.5791 \times 10^8$ |
| 2 | $I_{TA}^1$ | $3.1708 \times 10^7$ | $I_{TA}^2$ | $6.2339 \times 10^7$ | $I_{TA}^3$ | $1.4865 \times 10^8$ |
| 3 | $I_{TA}^1$ | $1.5584 \times 10^7$ | $I_{TA}^2$ | $3.0333 \times 10^7$ | $I_{TA}^3$ | $6.6731 \times 10^7$ |

**Figure 10.**
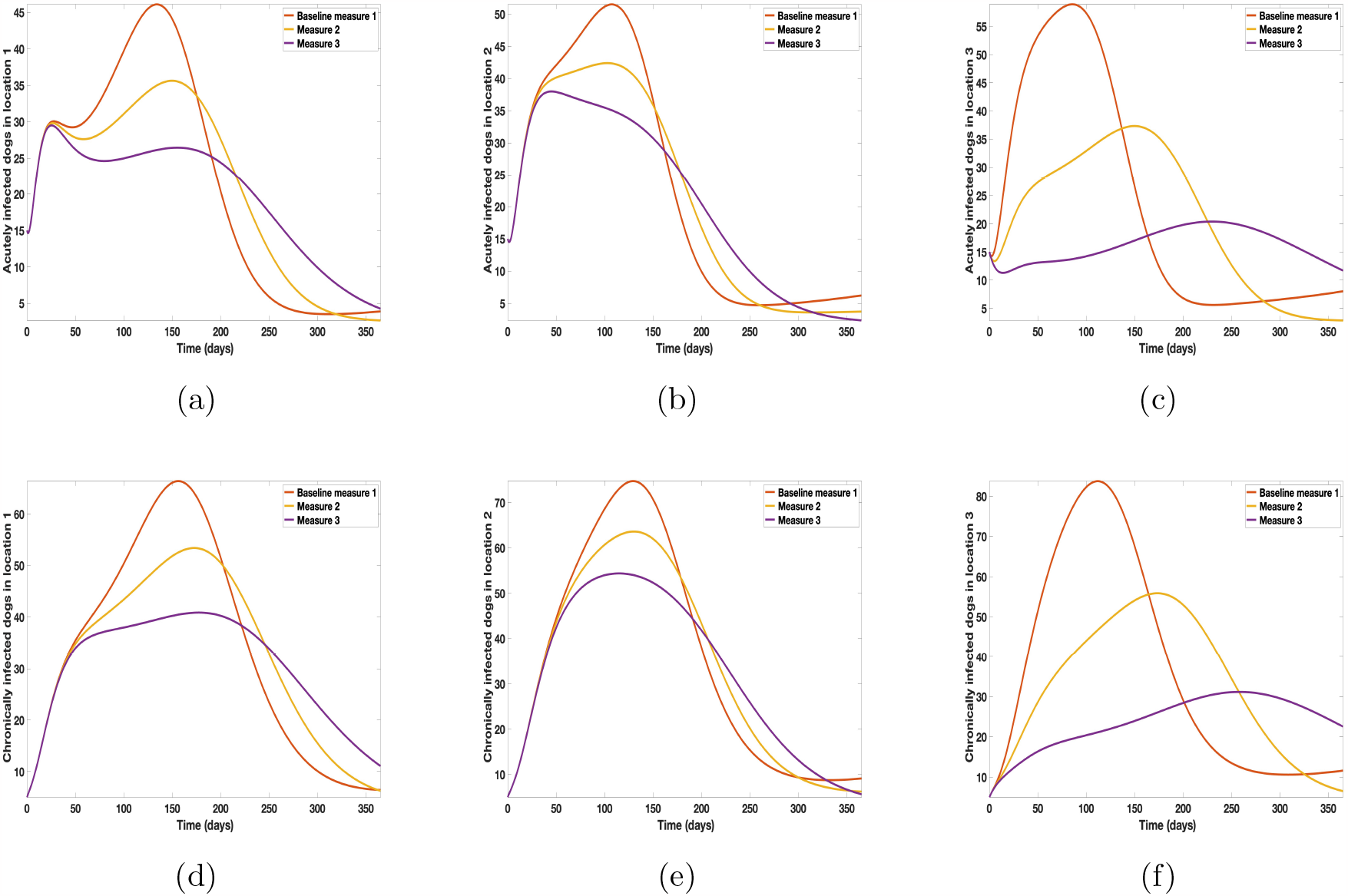
Simulation results of model (7) with movement. Control measures are only implemented in location 3. (a) acutely infected dogs; (b) chronically infected dogs; (c) infected larvae (d) infected nymphs (e) infected adult ticks.

### Connected locations: Control at all locations at the same time

With this scenario the three control measures described above are implemented in all three locations at the same time. We observed in all three locations significant reduction in population of acutely and chronically infected dogs using the three control measures, Measure 3 produced the most reduction see Figure 11. It is interesting to note that since similar levels of control are implemented in all three locations the effect of movement is cancelled out, as the sum of infected under this scenario is the same as the case with no movement. Similar trends are seen in Table 8 in the sum of acutely and chronically infected dogs and ticks over the simulation period. The outcome for infected larvae, nymphs, and adult ticks in each locations are shown in Figure F-5 in Appendix G.3.

**Table 8:**
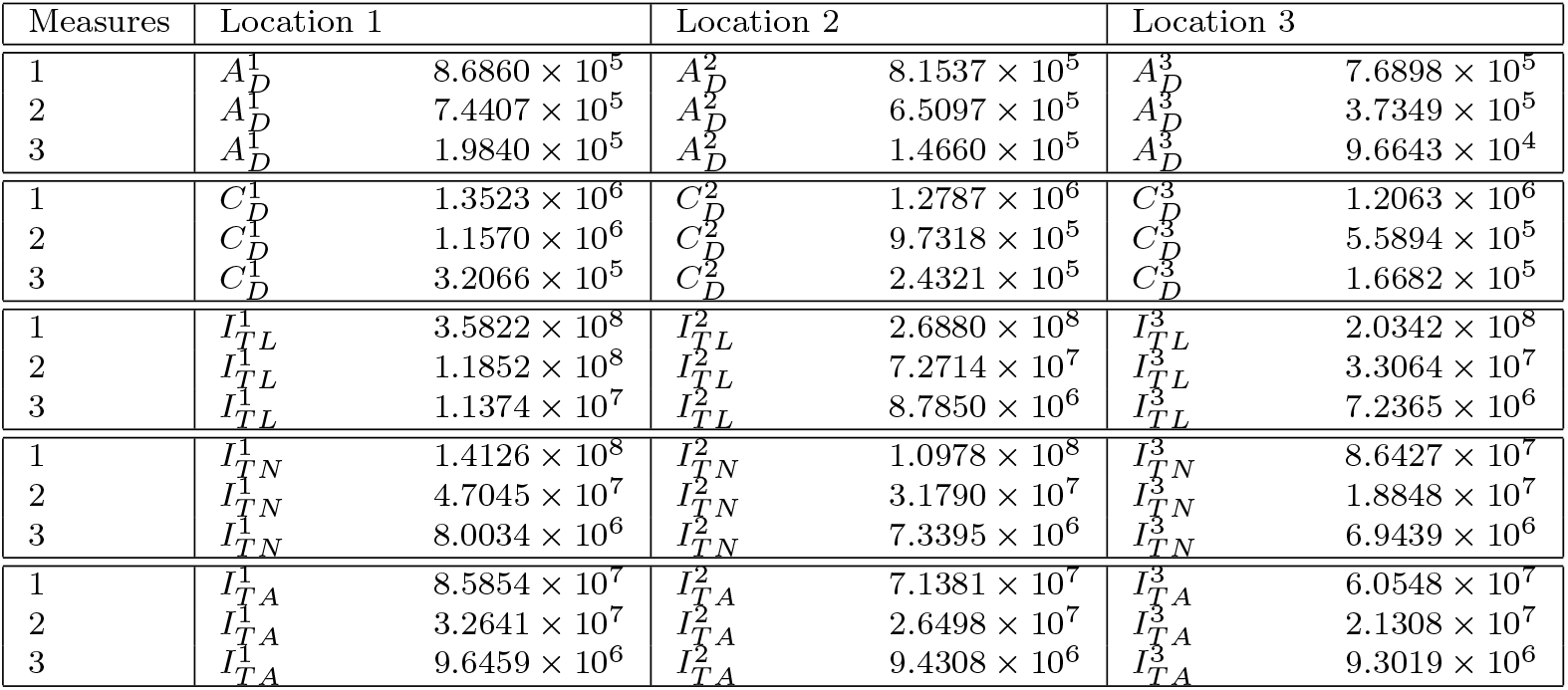
Sum of the simulation results of model (7) with movement. Control measures are implemented in all three locations 1, 2, and 3.

**Figure 11.**
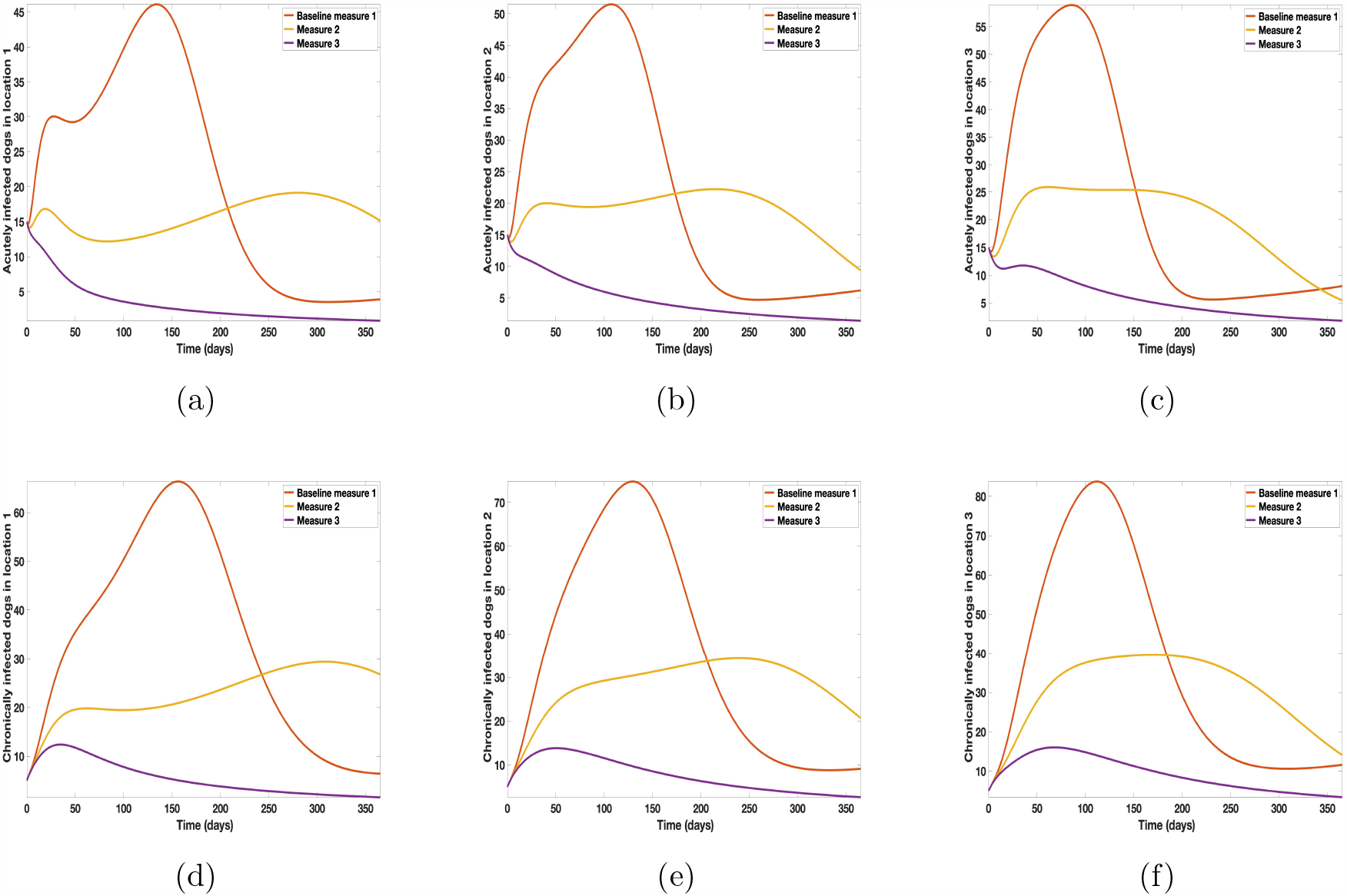
Simulation results of model (7) with movement. Control measures are implemented in all the three locations 1, 2, and 3. (a) acutely infected dogs; (b) chronically infected dogs; (c) infected larvae (d) infected nymphs (e) infected adult ticks.

## 4 Discussion and Conclusion

### Discussion

In this paper we developed and analyzed a mathematical model (6) for the disease transmission dynamics of *ehrlichia chaffeensis* in dogs using the natural history of infection of the disease. Features of this model include the different life stages of ticks and the different infectious stages of both dogs and ticks. The life stages of the ticks included egg, larvae, nymph, and adult which typically takes over a 2 year span to complete. The infectious stages of the dogs were divided into acute and clinical/chronic. The first 2-4 weeks of infection is referred to as the acute stage. After the initial 2-4 weeks if not treated the dog will progress into a clinical/chronic stage. This stage typically leads to death [21]. *Ehrlichia chaffeensis* is a serious tick-borne infectious disease that can cause life-threatening complications. It is important to know how to prevent dogs from being infected with *ehrlichia chaffeensis* and to be aware of the symptoms if dog becomes infected. One primary vector of *ehrlichia chaffeensis* is the *amblyomma americanum* tick.

We extend the *ehrlichia chaffeensis* model (6) by incorporating visitation and long distance migratory movements for dogs and ticks. Owners typically take their dogs on hikes or to dog parks. It is estimated that people take their dogs on a walk about 9 times a week for around 34 minutes. This usually ends up being a 2 mile walk. This totals to about 370 miles a year [36]. The short dog movements were captured through the visitation parameters *p*_*ij*_, which is the proportion of time a dog in location *i* spends visiting location *j*. Ticks long distance migratory movement may be due to ticks dropping off after feeding on either migratory birds moving north or from white-tail deer or other larger mammals [17].

Quantitative analysis of models (6) and (7) indicate that the disease-free equilibrium of these models are locally asymptotically stable when their reproduction number is less than one. Following [40], we can show that the global reproduction number of model (7) is bounded below and above by the local reproduction numbers from the patch with least and most transmission dynamics.

Next we carried out a global sensitivity analysis using LHS/PRCC method to determine the parameter with the most influence on the response functions sum of acutely and chronically infected (*A*_*D*_ + *C*_*D*_) and the sum of infected ticks in all life stages (*I*_*T L*_ +*I*_*T N*_ +*I*_*T A*_). To implement the analysis, we used both models (6) and (7), and parameter values obtained from literature in most cases as well as assumed some where we could not find their values. The significant parameters for model (7) with movement between the locations are disease progression rate in dogs (*σ*_*D*_), the rate (*υ*_*D*_) dogs progress to chronic infection from acute infection, dog recovery rate (*γ*_*D*_), the natural death rate of dogs (*μ*_*D*_), death rate (*δ*_*D*_) of acutely infected dogs, death rate of the chronically infected dogs (*δ*_*C*_), birth rate of dogs 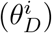, the maturation rates eggs to larvae (*α*_*E*_), the tick biting rate 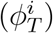, the dog transmission probability (*β*_*D*_), the tick transmission probability 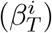 and death rate of ticks 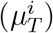, and the carrying capacity *K*^*i*^. There parameters are also significant when we used model (6) that models transmission dynamics in a single location, see Figure 2.

Furthermore, we observed in Figures 3 that the PRCC values of parameters 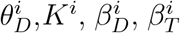, and 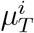 are relatively the same when there are no movement between the region. However, with movement the PRCC values for parameters 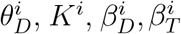, and 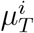 are different, with those in location 1 having higher values than those in locations 2 and 3 since there is more movement into location 1, than 2 and 3, see Figure 4. Knowing these significant parameters is essential to the formulation of effective control strategies for combating the spread of disease. For instance, a control that aims for a 10% decrease in the transmission probability in dogs (*β*_*D*_) will lead to 88% reduction in sume of acutely and chronically infected dogs and about 78% reduction in infected ticks of all life stages. Similarly a 10% decrease in the ticks transmission probability (*β*_*T*_) will lead to 71.4% reduced infection in dogs and about 83% reduced infection in ticks. Also, a 10% deduction in tick biting rate (*ϕ*_*T*_) will lead to about 95% reduction in infected dogs, and about 94% reduction in infected ticks. Furthermore, a 10% increase in ticks death rate (*μ*_*T*_) will result in about 72.4% reduction in infected dogs and about 83.4% decrease in ticks.

The simulation results of model (7) in Figure 5 show that locations with high infection rates like location 1 have a high number of infected dogs and ticks. This is due to the fact that the infected dogs and ticks are not moving the disease to other locations and keeping it localized in their location. On the other hand, locations with high movement and visitation rates see an increase in infected dogs and ticks. This is due to infected dogs and ticks traveling and infecting the dogs and ticks within the location that they are visiting (in the case of dogs) and moving to (in the case of ticks). In Figures 6(a) and 6(b) we see higher number of acutely and chronically infected dogs in location 3 followed by location 2 then location 1 even though infection is higher in location 1. Similar result is observed for infected larvae, nymphs adult ticks in Figures 6(c), 6(d), and 6(e). These results shows the impact of movement on the transmission of the disease as dogs and ticks move across locations. This result align with results in [28, 40], where the patch with the highest host movement or migration have the most infection. For instance Nguyen *et al*. [28] showed that deer mobility from a Lyme disease endemic county into Lyme disease free county will lead to the emergence of the disease in this second county free of the disease. Similarly, Zhang *et al*. [40] showed that rodent migration between patches can promote the disease spreading within all patches.

Next, we use the results from the sensitivity analysis coupled with movement between the locations to determine which control measure that reduces the most the spread of *ehrlichia chaffeensis* among dogs and ticks. Identifying these measures are crucial to decreasing the spread of *ehrlichia chaffeensis* amongst dogs. We note that during the single location control, the effect trickles to other locations due to the effect of movement between the locations. To see the impact of this single location control trickling effect, we compare in location 2 the effect of the control measures in location 1 only where the transmission is highest to the control measures in location 3 only where movement into it is highest, but transmission is lowest, we observed that controlling in location 3 only produces the most reduction in location 2. For instance, Measure 3 for acutely infected dogs in location 2 with Location 1 only control is 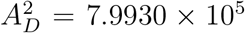, while acutely infected dogs in location 2 with location 3 only control is 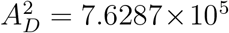. Similarly for the chronically infected dogs, we have in location 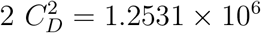 with location 1 only control, while 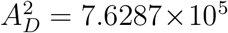with location 3 only control.

See Tables 5 and 7 for the other dogs and ticks variables.

### Conclusion

To conclude, the goal of this study was to develop a deterministic model of ordinary differential equations to gain insight in the transmission dynamics of *erhlichia chafeensis* between dogs and *amblyomma americanum* ticks. We found that infection in dogs and ticks are localized in the absence of movement and spreads between locations with highest infection in locations with the highest rate movement. We summarize the other results as follows:

i. The sensitivity analysis indicates that the on the sum of acutely and chronically infected (*A*_*D*_ + *C*_*D*_) and the sum of infected ticks in all life stages (*I*_*T L*_ + *I*_*T N*_ + *I*_*T A*_) with and without movement between the locations are disease progression rate in dogs, progression rate to chronic infection in dogs, dog recovery rate, the natural death rate of dogs, disease induced death rate (i) acutely and chronically infected dogs, birth rate of dogs, eggs maturation rates, tick biting rate, transmission probabilities in dogs and ticks, death rate of ticks, and the location carrying capacity.
ii. In the absence of movement, locations with high infection rates have a high number of infected dogs and ticks; while locations with high movement rates see an increase in infected dogs and ticks
iii. In a the single location control, the effect of the control measures which reduces infection trickles to other locations due to the effect of movement between the locations. Furthermore, most infection reduction trickling effect is observed in location with the most movement.

Thus, the results from this study show that dog owners should be careful about where their dogs visit, this will reduce *ehrlichia chaffeensis* in dogs; furthermore, they need to take their dogs to the vet immediately if any symptoms appear.

## Acknowledgments

This project is supported by the National Science Foundation under the EPSCOR Track 2 grant number 1920946.

## A Proof of Lemma 1

### Lemma 1

Let the initial data *F* (0) *≥* 0, where *F* (*t*) = (*S*_*D*_(*t*), *E*_*D*_(*t*), *A*_*D*_(*t*), *C*_*D*_(*t*), *R*_*D*_(*t*), *S*_*T E*_(*t*), *S*_*T L*_(*t*), *I*_*T L*_(*t*), *S*_*T N*_ (*t*), *I*_*T N*_ (*t*), *S*_*T A*_(*t*), *I*_*T A*_(*t*)). Then the solutions *F* (*t*) of the *ehrlichia chaffeensis* model (6) are non-negative for all *t >* 0. Furthermore

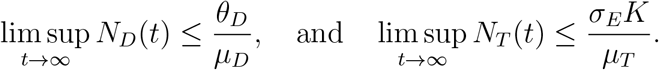

where

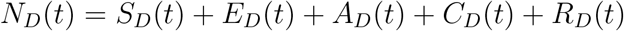

and

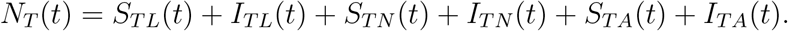

*Proof*. Let *t*_1_ = sup{*t >* 0 : *F* (*t*) *>* 0 *∈* [0, *t*]}. Thus, *t*_1_ *>* 0. It follows from the first equation of the system (6), that

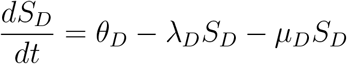

which can be re-written as

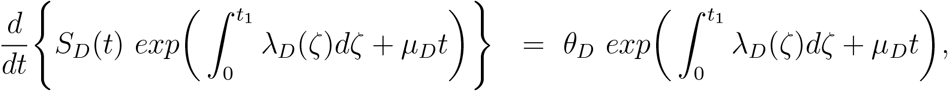

Hence,

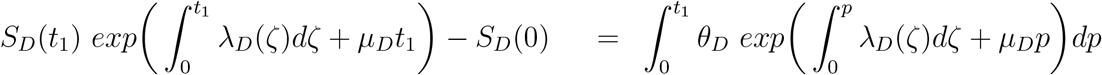

so that,

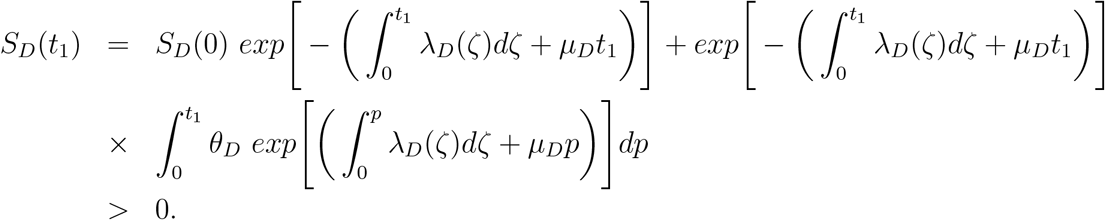

Similarly, it can be shown that *F >* 0 for all *t >* 0.

For the second part of the proof, note that 0 *< S*_*D*_(0) *≤ N*_*D*_(*t*), 0 *≤ E*_*D*_(0) *≤ N*_*D*_(*t*), 0 *≤ A*_*D*_(0) *≤ N*_*D*_(*t*), 0 *< C*_*D*_(0) *≤ N*_*D*_(*t*), 0 *≤ R*_*D*_(0) *≤ N*_*D*_(*t*), 0 *< S*_*T E*_(0) *≤ K*, 0 *< S*_*T L*_(0) *≤ N*_*T*_ (*t*), 0 *≤ I*_*T L*_(0) *≤ N*_*T*_ (*t*), 0 *< S*_*T N*_ (0) *≤ N*_*T*_ (*t*), 0 *≤ I*_*T N*_ (0) *≤ N*_*T*_ (*t*), 0 *< S*_*T A*_(0) *≤ N*_*T*_ (*t*), 0 *≤ I*_*T A*_(0) *≤ N*_*T*_ (*t*).

Adding the dog and tick component of the *ehrlichia chaffeensis* model (6) gives

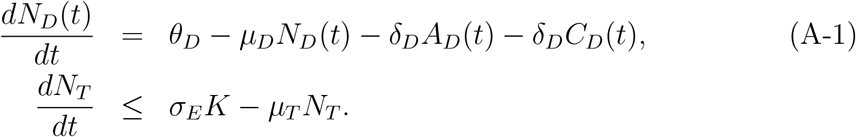

We suppose that *S*_*E*_ *< K*, where *K* is the carrying capacity.

Hence,

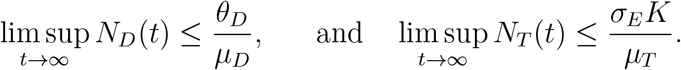

as required. □

## B Proof of Lemma 2

### Lemma 2.

The region 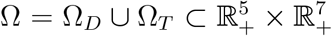 is positively-invariant for the model (6) with non-negative initial conditions in 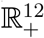.

*Proof*. It follows from the sum of the first five equations of model (6) that

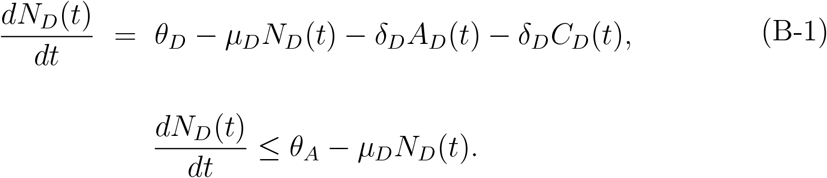

Hence, 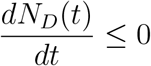, if 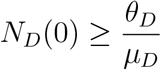. Thus,

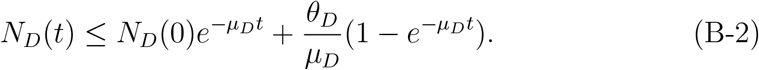

In particular, if 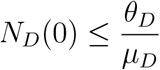, then 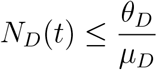.

Next, the last seven equations of model (6) give the following after summing the equations representing the larvae, nymphs, and adult stages

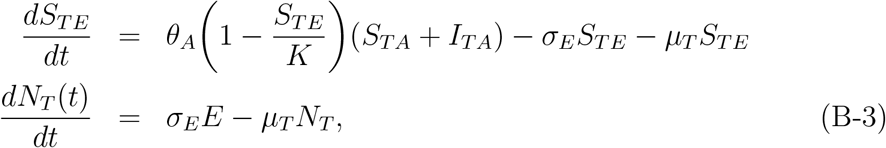

where *μ*_*T*_ = min{*μ*_*L*_, *μ*_*N*_, *μ*_*A*_}. Since *K* is the carrying capacity, it follows that *S*_*T E*_ *≤ K*. Hence, equation (B-3) becomes

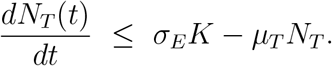

Thus,

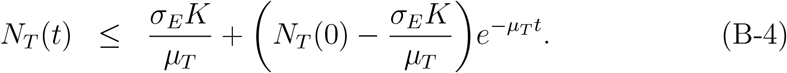

Furthermore, if 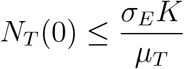, then 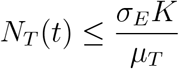.

Equations (B-2), and (B-4) implies that *N*_*D*_(*t*), and *N*_*T*_ (*t*) are bounded and all solutions starting in the region Ω remain in Ω. Thus, the region is positively-invariant and hence, the region Ω attracts all solutions in 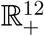. □

## C Basic qualitative properties of model (7)

### C.1 Positivity and boundedness of solutions

For the *ehrlichia chaffeensis* transmission model (7) to be epidemiologically meaningful, it is important to prove that all its state variables are non-negative for all time. In other words, solutions of the model system (7) with non-negative initial data will remain non-negative for all time *t >* 0.

#### Lemma 4.

*Let the initial data F* (0) *≥* 0, *where* 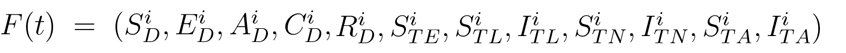. *Then the solutions F* (*t*) *of the ehrlichia chaffeensis model* (7) *are non-negative for all t >* 0. *Furthermore*

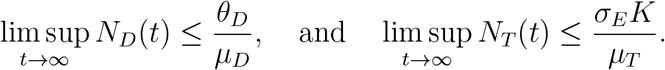

*where* 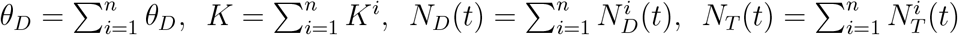 *with* 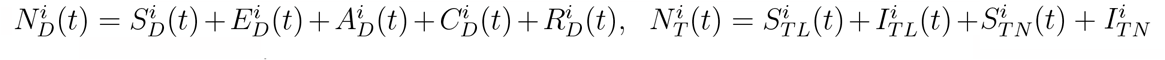.

*Proof*. Let *t*_1_ = sup{*t >* 0 : *F* (*t*) *>* 0 *∈* [0, *t*]}. Thus, *t*_1_ *>* 0. It follows from the seventh equation of the system (7), that

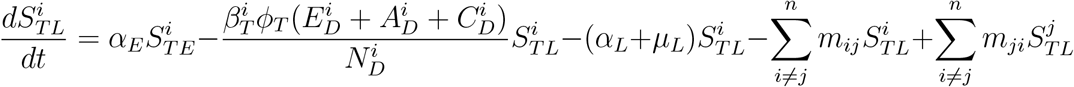

which can be re-written as

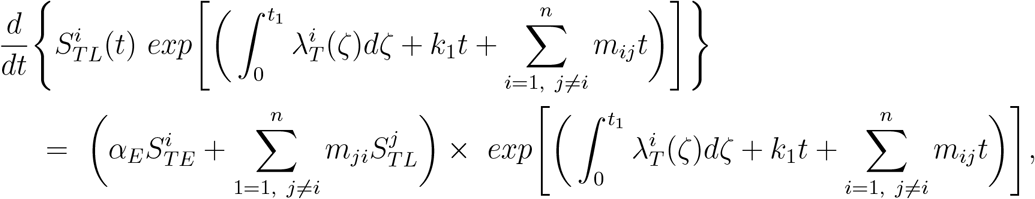

where 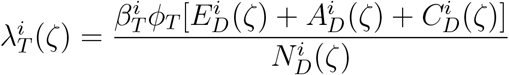, and *k*_1_ = *α*_*L*_ + *μ*_*L*_. Hence,

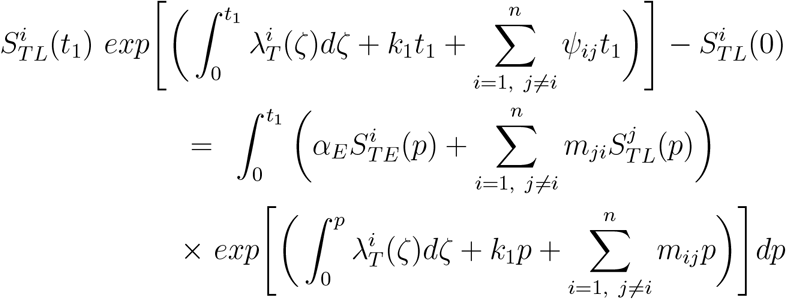

so that,

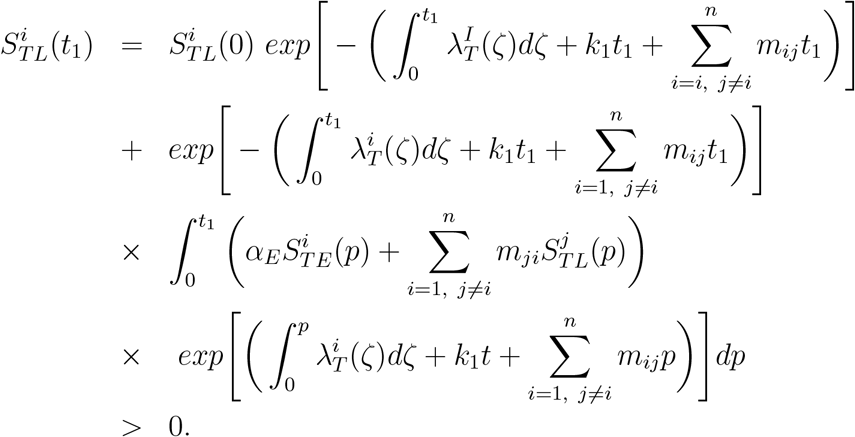

Similarly, it can be shown that *F >* 0 for all *t >* 0.

For the second part of the proof, note that 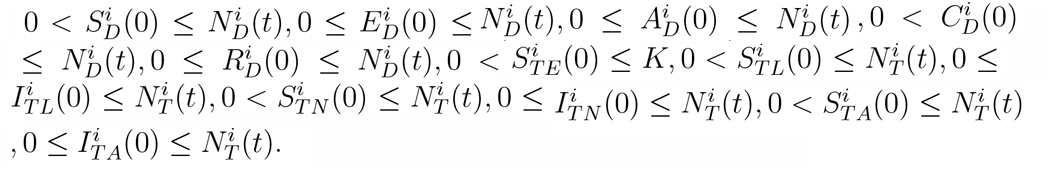.

Considering the dog and tick components of model (6) we have

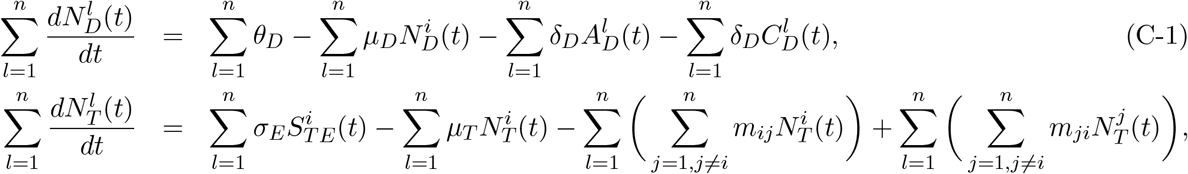

where. 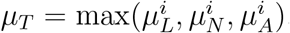. Since 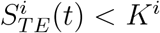, with *K*^*i*^ being the carrying capacity of each of region *i*, we have

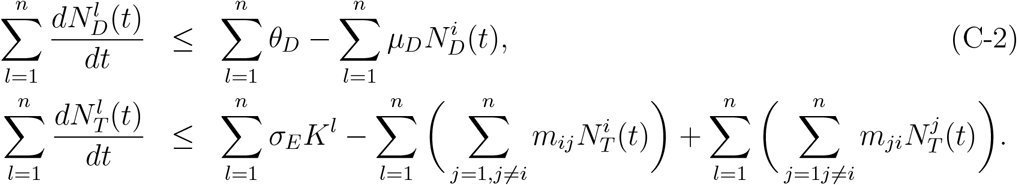

Now summing separately the dog and tick components, gives

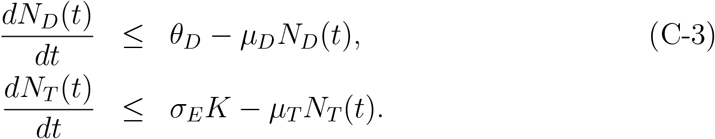

where 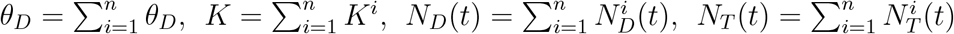 with 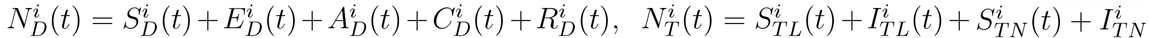.Also,

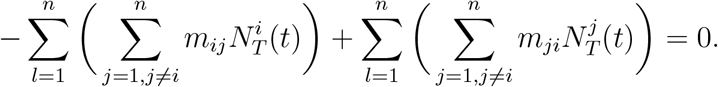

Thus, equation (C-3) lead to the following

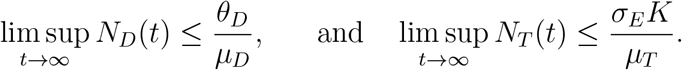

as required. □

### C.2 Invariant regions

The *ehrlichia chaffeensis* model (7) will be analyzed in a biologically-feasible region as follows. Consider the feasible region

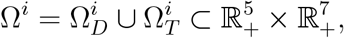

where,

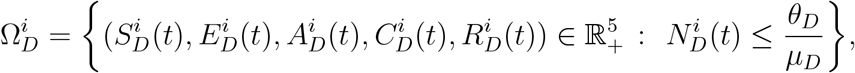

and

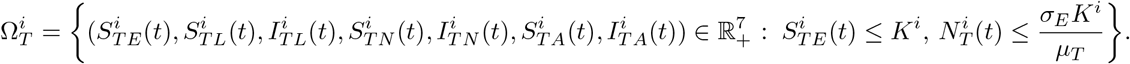

with 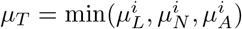.

#### Lemma 5.

*The region* 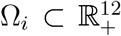 *is positively-invariant for the ehrlichia chaffeensis model* (7) *with non-negative initial conditions in* 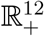.

*Proof*. The following steps are followed to establish the positive invariance of Ω^*i*^ (i.e., solutions in Ω^*i*^ remain in Ω^*i*^ for all *t >* 0). The rate of change of the total dog and tick populations is obtained by adding separately the dog and tick component of model (7) for a particular region

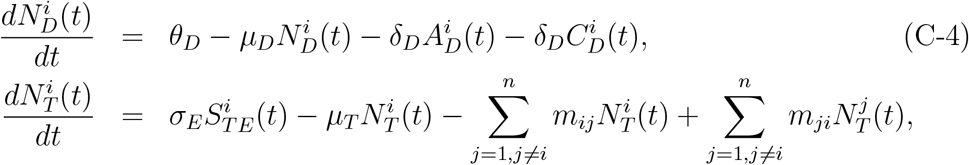

where 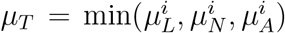. Since 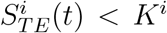, with *K*^*i*^ being the carrying capacity of each of region *i*, equation (C-4) reduces to

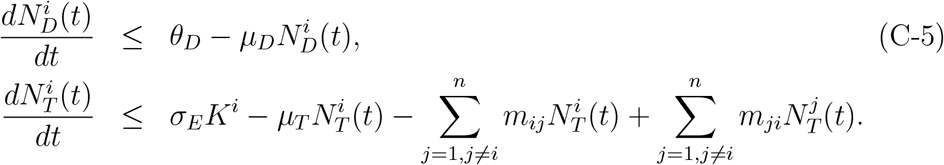

From (C-5), and using a standard comparison theorem [22], we can show that

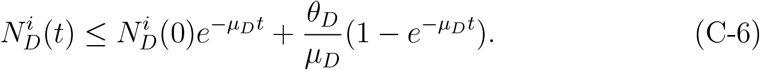

and, 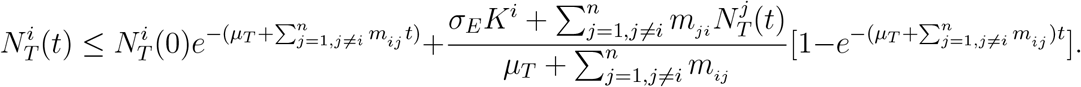 Furthermore, if 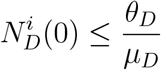, then 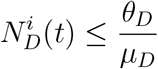 and 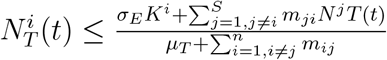 if 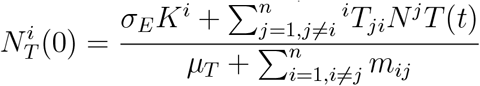.

Thus, the region Ω is positively-invariant. Hence, it is sufficient to consider the dynamics of the flow generated by (7) in Ω^*i*^. In this region, the model is epidemiological and mathematically well-posed [18]. Thus, every solution of the basic model (7) with initial conditions in Ω remains in Ω for all *t >* 0. Therefore, the *ω*-limit sets of the system (7) are contained in Ω^*i*^. This result is summarized below.

## D Stability of disease-free equilibrium (DFE)

In the next section, the conditions for the existence and stability of the disease-free equilibrium of the model *ehrlichia chaffeensis* (7) are stated.

The disease-free equilibrium of the *ehrlichia chaffeensis* model (7) with movement DFE is given by

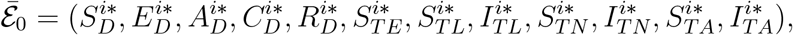

where,

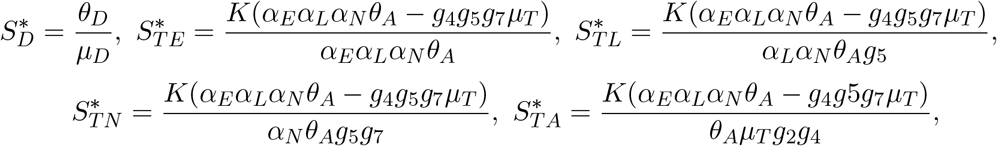

The stability of *ε* _0_ can be established using the next generation matrix method on system (7). Taking *E*_*D*_, *A*_*D*_, *C*_*D*_, *I*_*T L*_, *I*_*T N*_, and *I*_*T A*_ as the infected compartments and then using the aforementioned notation, the Jacobian 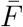 and 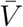 matrices for new infectious terms and the remaining transfer terms, respectively, are defined as:

## E Stability of disease-free equilibrium (DFE)

Therefore, the reproduction is given as 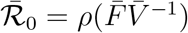, where

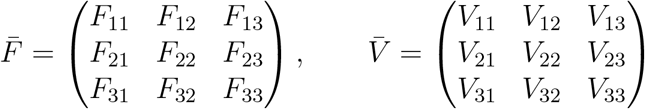

with

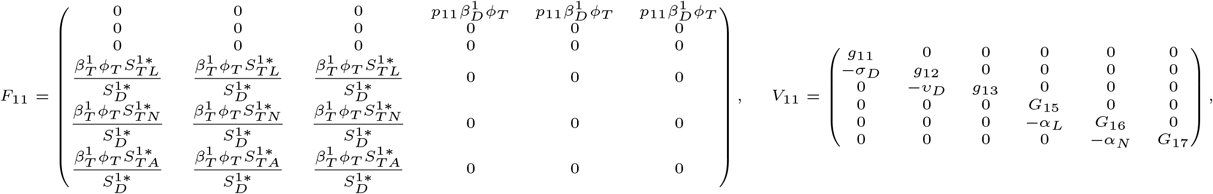

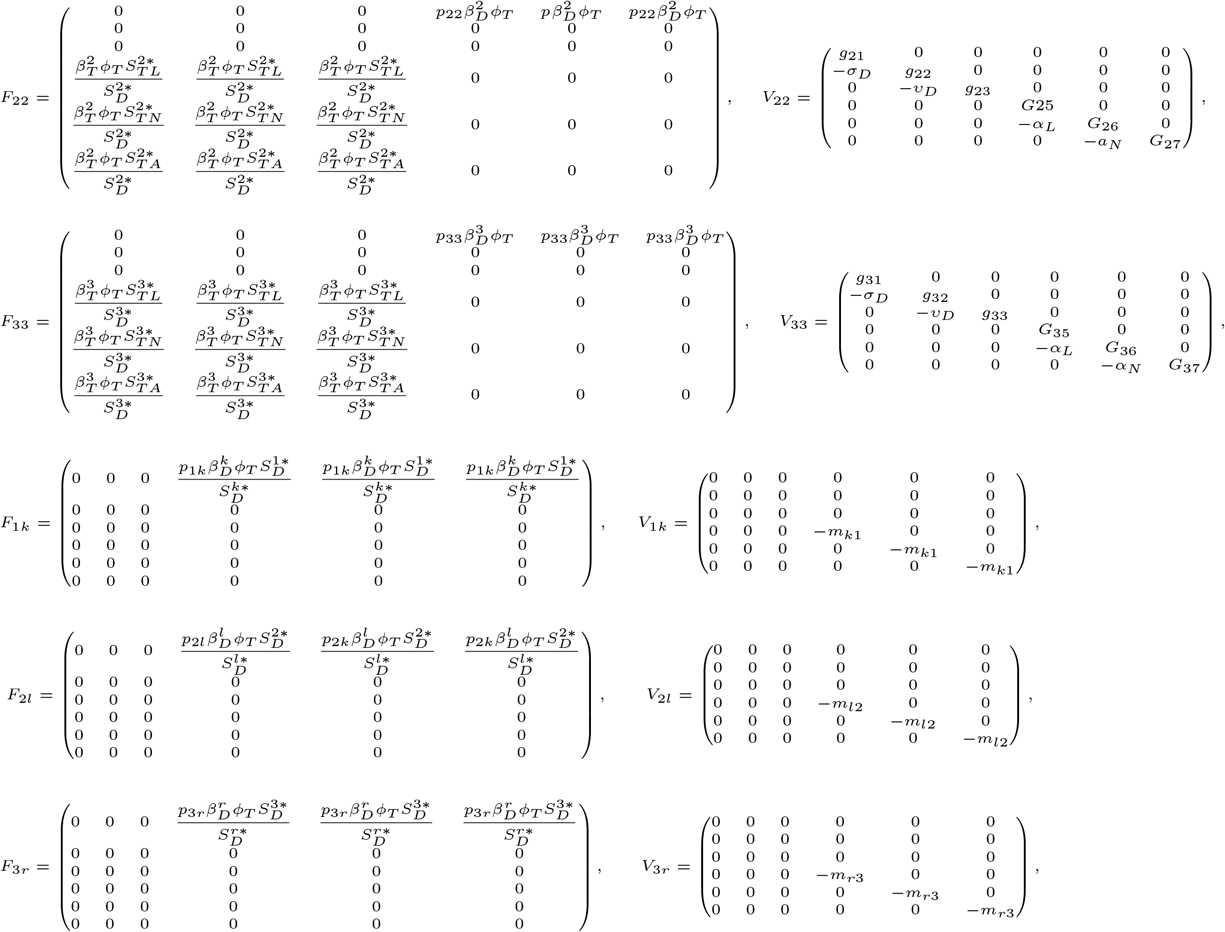

and 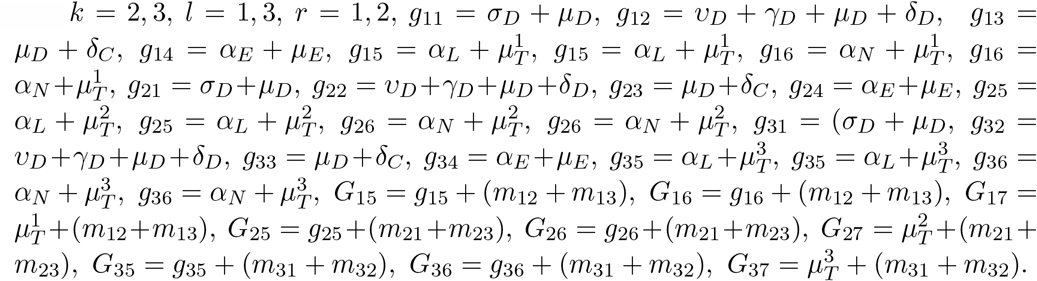

The following result is established using Theorem 2 in [39].

### Lemma 6.

*The DFE of the ehrlichia chaffeensis model* (7) *with movement, given by* 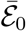, *is locally asymptotically stable (LAS) if* 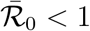, *and unstable if* 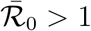.

The basic reproduction number 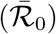 measures the average number of new infections generated by a single infected individual (tick or dog) in a completely susceptible population [4, 15, 18, 39]. Thus, Lemma 6 implies that malaria can be eliminated from the human population (when 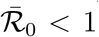) if the initial sizes of the sub-populations are in the basin of attraction of the DFE, 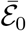.

## F Sensitivity analysis with movement

Table E-1 give the result of the PRCC and p-values of the *ehrlichia chaffeensis* model (6) when the locations are isolated and there are no movement between them.

**Table E-1:** PRCC and p-values of the *ehrlichia chaffeensis* model (7) with no movement between the regions using as response functions the sum of infected dogs (*A*_*D*_ + *C*_*D*_) and the sum of infected tick (*I*_*T L*_+*I*_*T N*_ +*I*_*T A*_) with parameter values in Table 2 that are with *±*20% ranges from the baseline values. With this movement scenario, the locations are isolated with no movement between them.

| Parameters | $A_D + C_D$ | | $I_{TL} + I_{TN} + I_{TA}$ | |
| --- | --- | --- | --- | --- |
|  | PRCC | p-value | PRCC | p-value |
| $\varepsilon$ | 0.0279 | 0.3853 | 0.0378 | 0.2388 |
| $\sigma_D$ | -0.5088 | 0.0000 | -0.6808 | 0.0000 |
| $\nu_D$ | 0.6599 | 0.0000 | 0.3462 | 0.0000 |
| $\gamma_D$ | -0.7750 | 0.0000 | -0.4917 | 0.0000 |
| $\mu_D$ | 0.0037 | 0.9094 | 0.0704 | 0.0281 |
| $\delta_D$ | -0.6603 | 0.0000 | -0.2715 | 0.0000 |
| $\delta_C$ | -0.8769 | 0.0000 | -0.5871 | 0.0000 |
| $\theta_A$ | 0.0156 | 0.6265 | -0.0128 | 0.6897 |
| $\mu_E$ | -0.0048 | 0.8816 | -0.0038 | 0.9058 |
| $\alpha_E$ | 0.8402 | 0.0000 | 0.9183 | 0 |
| $\alpha_L$ | 0.0032 | 0.9202 | -0.0270 | 0.4003 |
| $\alpha_N$ | 0.0670 | 0.0365 | 0.0655 | 0.0410 |
| $\phi_T$ | 0.9747 | 0 | 0.9750 | 0 |
| $\theta_D^3$ | -0.0008 | 0.9793 | -0.0441 | 0.1698 |
| $\beta_D^3$ | 0.6703 | 0.0000 | 0.5685 | 0.0000 |
| $\beta_T^3$ | 0.4351 | 0.0000 | 0.5932 | 0.0000 |
| $\mu_T^3$ | -0.4789 | 0.0000 | -0.6239 | 0.0000 |
| $K^3$ | 0.4471 | 0.0000 | 0.5875 | 0.0000 |
| $\theta_D^2$ | 0.0169 | 0.5978 | -0.0382 | 0.2340 |
| $\beta_D^2$ | 0.6680 | 0.0000 | 0.5452 | 0.0000 |
| $\beta_T^2$ | 0.4263 | 0.0000 | 0.5951 | 0.0000 |
| $\mu_T^2$ | -0.4809 | 0.0000 | -0.6248 | 0.0000 |
| $K^2$ | 0.4538 | 0.0000 | 0.6034 | 0.0000 |
| $\theta_D^1$ | -0.0074 | 0.8167 | -0.0102 | 0.7496 |
| $\beta_D^1$ | 0.6688 | 0.0000 | 0.5217 | 0.0000 |
| $\beta_T^1$ | 0.4372 | 0.0000 | 0.6020 | 0.0000 |
| $\mu_T^1$ | -0.4968 | 0.0000 | -0.6358 | 0.0000 |
| $K^1$ | 0.4457 | 0.0000 | 0.6019 | 0.0000 |

Table E-2: gives the result of the PRCC and p-values of the *ehrlichia chaffeensis* model (6) when the locations are connected and there are movement of dogs and ticks between them.

**Table E-2:** PRCC and p-values of the *ehrlichia chaffeensis* model (7) with movement between the regions using as response functions the sum of infected dogs (*A*_*D*_ + *C*_*D*_) and the sum of infected tick (*I*_*T L*_ + *I*_*T N*_ + *I*_*T A*_) with parameter values in Table 2 that are with *±*20% ranges from the baseline values. With the movement scenario, the locations are connect.

| Parameters | $A_D + C_D$ | | $I_{TL} + I_{TN} + I_{TA}$ | |
| --- | --- | --- | --- | --- |
|  | PRCC | p-value | PRCC | p-value |
| $\epsilon$ | 0.0638 | 0.0468 | 0.0618 | 0.0539 |
| $\sigma_D$ | -0.4702 | 0.0000 | -0.7006 | 0.0000 |
| $v_D$ | 0.6496 | 0.0000 | 0.3912 | 0.0000 |
| $\gamma_D$ | -0.7530 | 0.0000 | -0.5108 | 0.0000 |
| $\mu_D$ | -0.0645 | 0.0444 | 0.0683 | 0.0331 |
| $\delta_D$ | -0.6403 | 0.0000 | -0.2912 | 0.0000 |
| $\delta_C$ | -0.8629 | 0.0000 | -0.5626 | 0.0000 |
| $\theta_A$ | 0.0283 | 0.3772 | 0.0023 | 0.9437 |
| $\mu_E$ | 0.0416 | 0.1947 | 0.0250 | 0.4360 |
| $\alpha_E$ | 0.8030 | 0.0000 | 0.9235 | 0 |
| $\alpha_L$ | -0.0527 | 0.1001 | -0.0071 | 0.8241 |
| $\alpha_N$ | 0.0036 | 0.9112 | -0.0035 | 0.9125 |
| $\phi_T$ | 0.9466 | 0 | 0.9695 | 0 |
| $\theta_D^3$ | -0.0070 | 0.8269 | -0.0396 | 0.2172 |
| $\beta_D^3$ | 0.3107 | 0.0000 | 0.1731 | 0.0000 |
| $\beta_T^3$ | 0.1252 | 0.0001 | 0.1774 | 0.0000 |
| $\mu_T^3$ | -0.0801 | 0.0125 | -0.1810 | 0.0000 |
| $K^3$ | 0.4308 | 0.0000 | 0.6394 | 0.0000 |
| $\theta_D^2$ | 0.0066 | 0.8367 | -0.0083 | 0.7971 |
| $\beta_D^2$ | 0.4950 | 0.0000 | 0.3786 | 0.0000 |
| $\beta_T^2$ | 0.2678 | 0.0000 | 0.4716 | 0.0000 |
| $\mu_T^2$ | -0.2749 | 0.0000 | -0.5064 | 0.0000 |
| $K^2$ | 0.4011 | 0.0000 | 0.6311 | 0.0000 |
| $\theta_D^1$ | 0.0476 | 0.1378 | 0.0014 | 0.9646 |
| $\beta_D^1$ | 0.4981 | 0.0000 | 0.6585 | 0.0000 |
| $\beta_T^1$ | 0.6794 | 0.0000 | 0.8616 | 0.0000 |
| $\mu_T^1$ | -0.5074 | 0.0000 | -0.7708 | 0.0000 |
| $K^1$ | 0.4292 | 0.0000 | 0.6408 | 0.0000 |

## G Simulating the *ehrlichia chaffeensis* model (7)

### G.1 Control in isolated locations

**Figure F-1:**
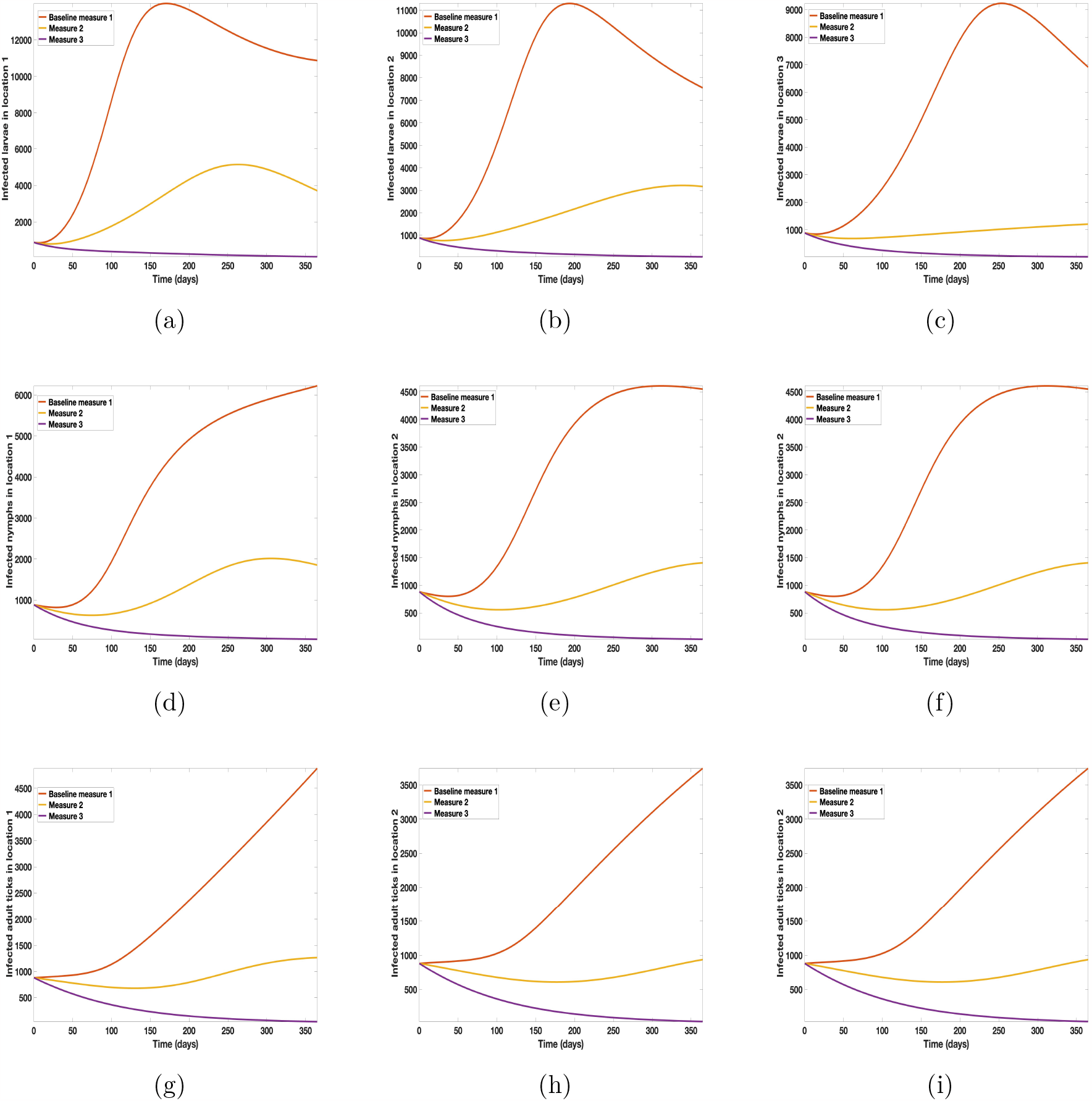
Simulation results of model (7) with no movement using parameters given in Table 2. Infection is higher in location 1 followed by location 2, location 3 has the least infection. More movement to location 3 from locations 1, and 3. (a) - (c) infected larvae (d) - (f) infected nymphs (g) - (i) infected adult ticks.

### G.2 Connected locations: Control in one patch at a time

#### G.2.1 Connected locations: Control in location 1 only

**Figure F-2:**
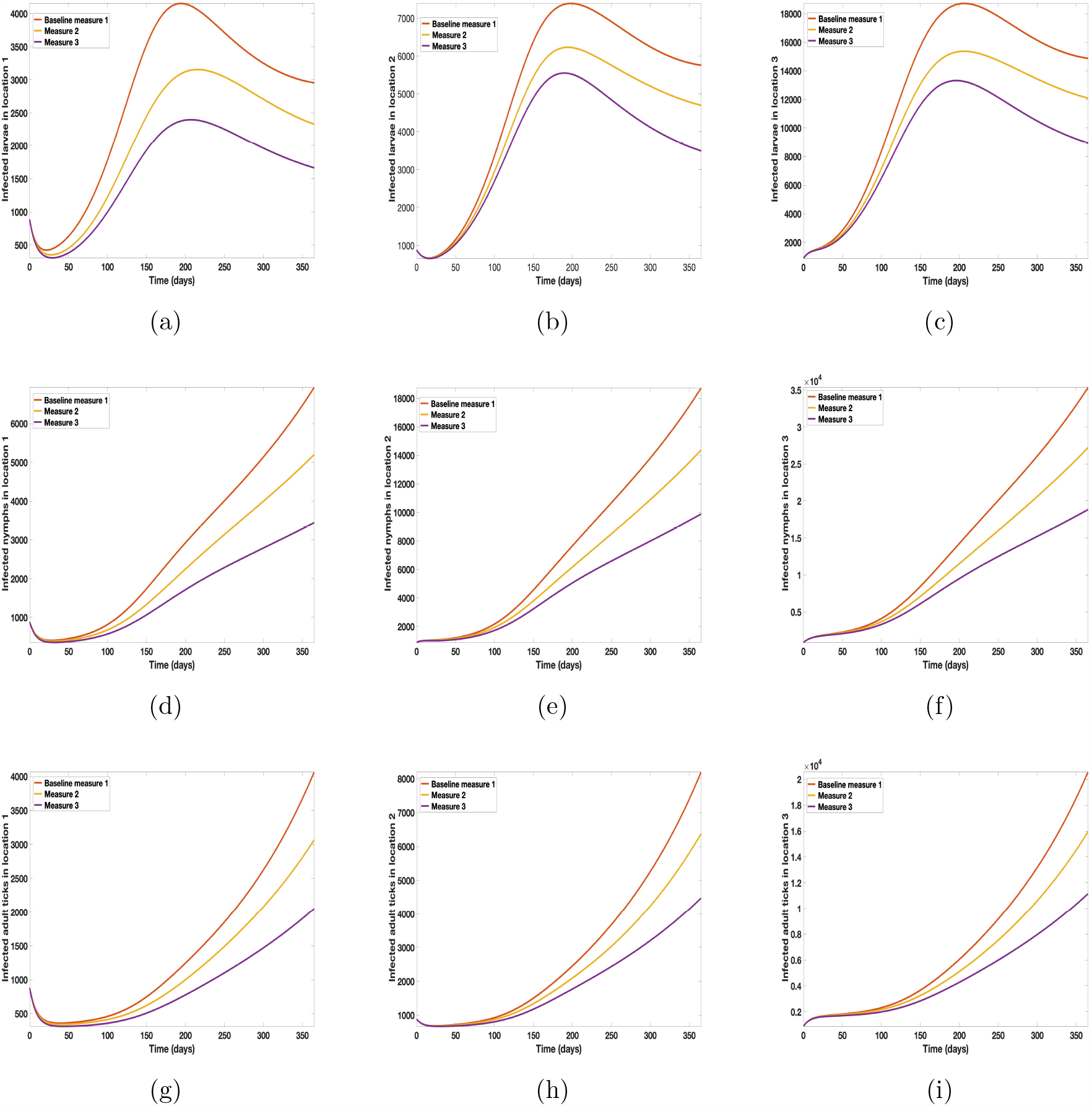
Simulation results of model (7) with movement. Control measures are implemented in location 1 only. (a) - (c) infected larvae (d) - (f) infected nymphs (g) - (i) infected adult ticks.

#### G.2.2 Connected locations: Control in location 2 only

**Figure F-3:**
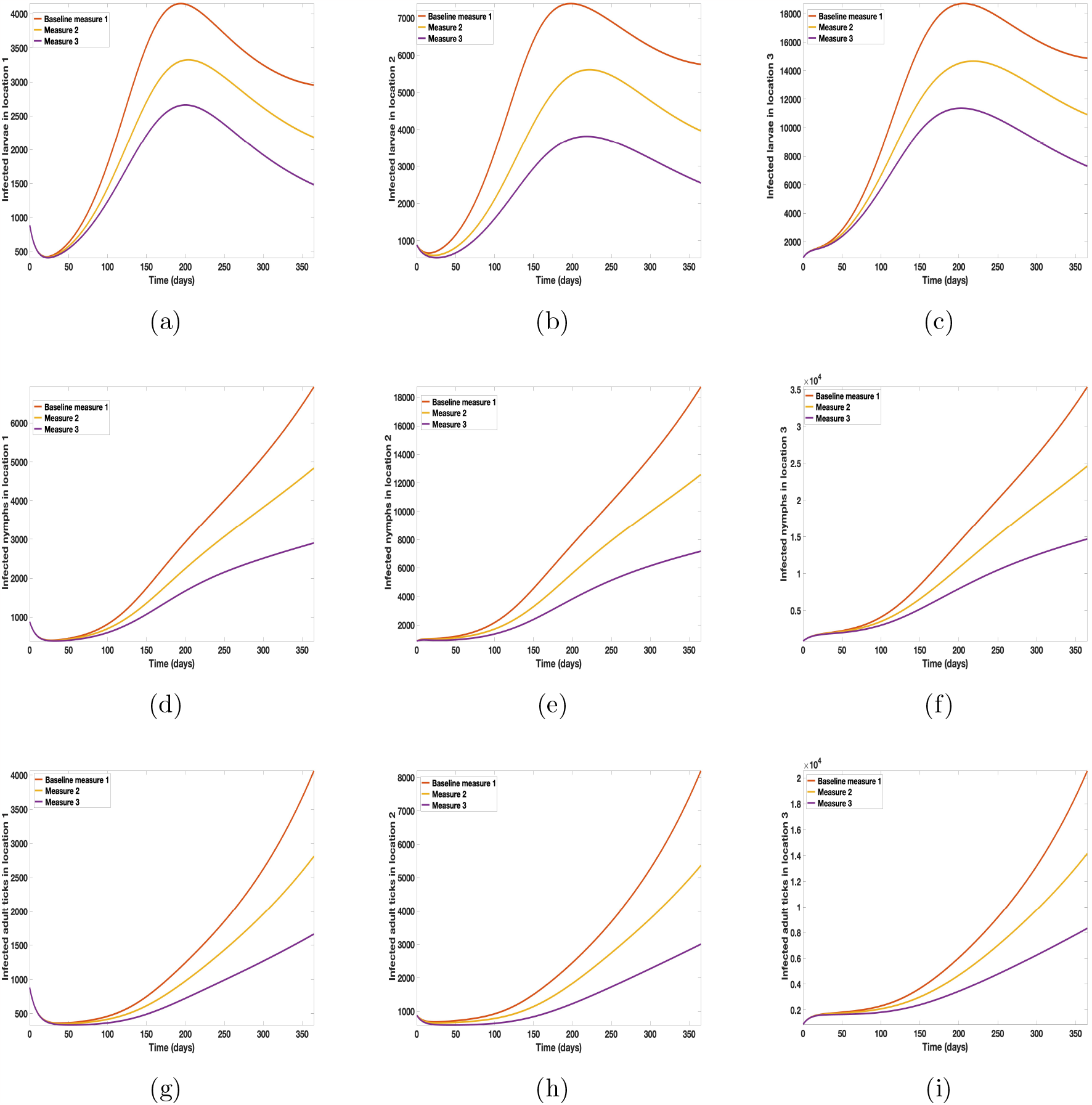
Simulation results of model (7) with movement. Control measures are implemented in location 2 only. (a) - (c) infected larvae (d) - (f) infected nymphs (g) - (i) infected adult ticks.

#### G.2.3 Connected locations: Control in location 3 only

**Figure F-4:**
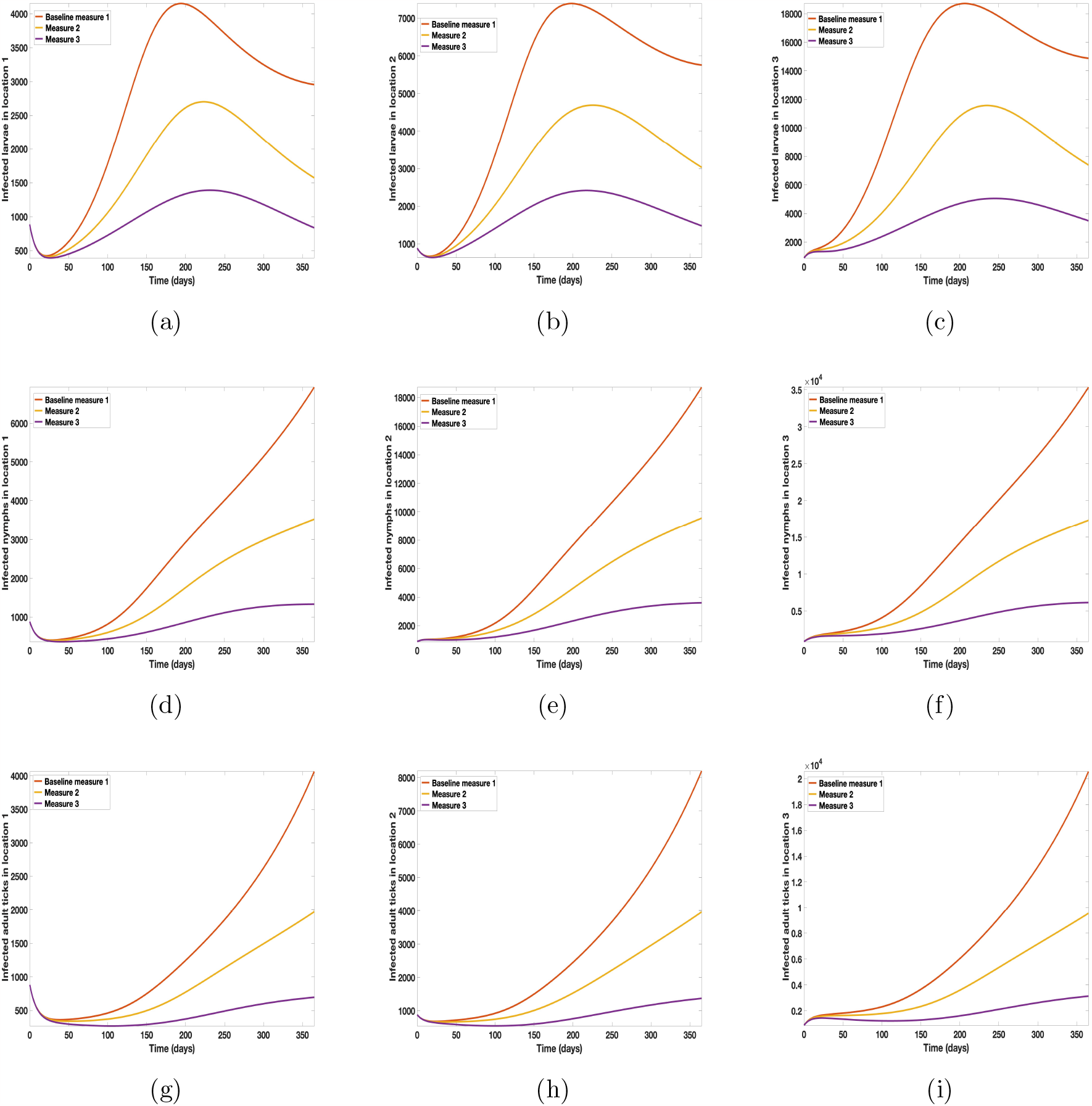
Simulation results of model (7) with movement. Control measures are implemented in location 3 only. (a) - (c) infected larvae (d) - (f) infected nymphs (g) - (i) infected adult ticks.

### G.3 Control in connected locations

**Figure F-5:**
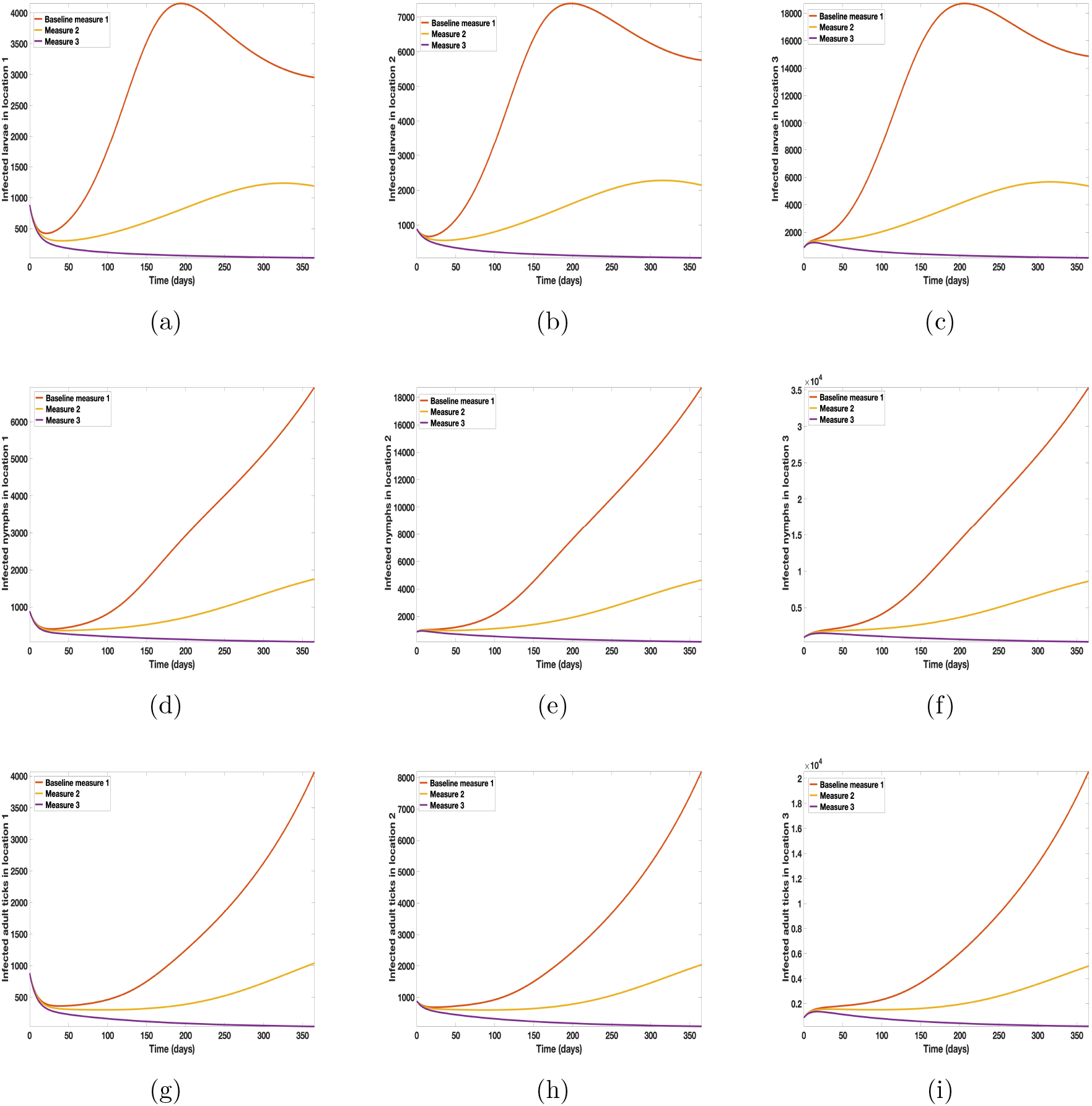
Simulation results of model (7) with movement. Control measures are implemented in all three locations 1, 2, and 3. (a) - (c) infected larvae (d) - (f) infected nymphs (g) - (i) infected adult ticks.

## References

[1] Folashade Agusto, Amy Goldberg, Omayra Ortega, Joan Ponce, Sofya Zaytseva, Suzanne Sindi, and Sally Blower. How do interventions impact malaria dynamics between neighboring countries? a case study with botswana and zimbabwe. Using Mathematics to Understand Biological Complexity: From Cells to Populations, pages 83–109, 2021.

[2] Folashade B Agusto. Malaria drug resistance: the impact of human movement and spatial heterogeneity. Bulletin of mathematical biology, 76(7):1607–1641, 2014.

[3] Folashade B Agusto and Soyeon Kim. Impact of mobility on methicillinresistant staphylococcus aureus among injection drug users. Antibiotics, 8(2):81, 2019.

[4] R. M. Anderson and R. May. Infectious Diseases of Humans. Oxford University Press, New York., 1991.

[5] Anna Burke. How long do dogs live? https://www.akc.org/expert-advice/health/how-long-do-dogs-live/, Accessed August 7,2022.

[6] GrahamJ. Hickling Antoinette Ludwig, Howard S. Ginsberg and Nicholas H. Ogden. A dynamic population model to investigate effects of climate and climate-independent factors on the lifecycle of amblyomma americanum. Journal of Medical Entomology, 2015. Accessed September 19 2022.

[7] Julien Arino, Jonathan R Davis, David Hartley, Richard Jordan, Joy M Miller, and P Van Den Driessche. A multi-species epidemic model with spatial dynamics. Mathematical Medicine and Biology, 22(2):129–142, 2005.

[8] R.B. Atherton, R.W. Schainker and E.R. Ducot. On the statistical sensitivity analysis of models for chemical kinetic. AIChE Journal, 21(3):441–448, 1975.

[9] S.M. Blower and H.I. Dowlatabadi. Sensitivity and uncertainty analysis of complex models of disease transmission: an hiv model, as an example. International Statistical Review/Revue Internationale de Statistique,pages 229–243, 1994.

[10] CDC. Ehrlichiosis in dogs: Fast facts for veterinarians. https://www.cdc.gov/ehrlichiosis/pdfs/fs-ehrlichiosisvet-508.pdf, Accessed January 16,2023.

[11] Center for Disease Prevention and Control (CDC). Transmission and epidemeology. https://www.cdc.gov/ehrlichiosis/healthcare-providers/transmission-and-epidemiology.html, 2019. Accessed August 7,2022.

[12] Chelsie Wheeler. The life cycle of a tick. https://study.com/learn/lesson/tick-life-cycle-reproduction-eggs.html, Accessed August 7 2022.

[13] Nakul Chitnis, James M Hyman, and Jim M Cushing. Determining important parameters in the spread of malaria through the sensitivity analysis of a mathematical model. Bulletin of mathematical biology, 70(5):1272–1296, 2008.

[14] Chris Cosner, John C Beier, Robert Stephen Cantrell, D Impoinvil, Lev Kapitanski, Matthew David Potts, A Troyo, and Shigui Ruan. The effects of human movement on the persistence of vector-borne diseases. Journal of theoretical biology, 258(4):550–560, 2009.

[15] O. Diekmann, J.A.P. Heesterbeek, and J.A.P. Metz. On the definition and computation of the basic reproduction ratio r0 in models for infectious diseases in heterogeneous populations. Journal of Mathematical Biology, 28:503–522, 1990.

[16] D.M. Hamby. A review of techniques for parameter sensitivity analysis of environmental models. Environmental monitoring and assessment, 32(2):135–154, 1994.

[17] Jane M Heffernan, Yijun Lou, and Jianhong Wu. Range expansion of ixodes scapularis ticks and of borrelia burgdorferi by migratory birds. Discrete Contin Dyn Syst Ser B, 19(10):3147–3167, 2014.

[18] H.W. Hethcote. The mathematics of infectious diseases. SIAM Review, 42(4):599–653, 2000.

[19] Indiana Department of Health. Amblyomma americanum. https://www.in.gov/health/erc/zoonotic-and-vectorborne-epidemiology-entomolpests/amblyomma-americanum/, 2022. Accessed August 7,2022.

[20] Jonathan Sellman, MD MPH. Vietnam war dogs and an unknown new disease. https://jonathansellmanmd.com/ x2015/06/17/420/, Accessed January 2,2023.

[21] Krista Williams, BSc, DVM, CCRP; Ryan Llera, BSc, DVM; Ernest Ward, DVM. Ehrlichiosis in dogs. https://vcahospitals.com/know-your-pet/ehrlichiosis-in-dogs, Accessed August 7, 2022.

[22] Vangipuram Lakshmikantham, Srinivasa Leela, and Anatoly A Martynyuk. Stability analysis of nonlinear systems. Springer, 1989.

[23] Lauren Leonardi. Identify ehrlichiosis symptoms at all 3 stages of the disease learn what signs to watch out for. https://www.petcarerx.com/article/identify-ehrlichiosis-symptoms-at-all-3-1545, 2021. Accessed August 7,2022.

[24] Milliward Maliyoni, Holly D Gaff, Keshlan S Govinder, and Faraimunashe Chirove. Multipatch stochastic epidemic model for the dynamics of a tick-borne disease. Frontiers in Applied Mathematics and Statistics, 9:1122410, 2023.

[25] Mar Vista Animal Medical Center. Ehrlichia infection (canine). https://www.marvistavet.com/ehrlichia-infection-canine.pml, 2020. Accessed August 7,2022.

[26] S. Marino, I.B. Hogue, C.J. Ray, and D. E. Kirschner. A methodology for performing global uncertainty and sensitivity analysis in systems biology. Journal of theoretical biology, 254(1):178–196, 2008.

[27] M.D. McKay, R.J. Beckman, and W.J. Conover. A comparison of three methods for selecting values of input variables in the analysis of output from a computer code. Technometrics, 42(1):55–61, 2000.

[28] Aileen Nguyen, Joseph Mahaffy, and Naveen K Vaidya. Modeling transmission dynamics of lyme disease: Multiple vectors, seasonality, and vector mobility. Infectious Disease Modelling, 4:28–43, 2019.

[29] NorthShoreAnimalLeague. Prevent a litter-spay and neuter your pets. https://www.animalleague.org/wp-content/uploads/2017/06/dogs-multiply-pyramid.pdf, Accessed August 7,2022.

[30] Shigui Ruan, Wendi Wang, Simon A Levin, et al. The effect of global travel on the spread of sars. Mathematical Biosciences and Engineering, 3(1):205, 2006.

[31] S. Harrus, P.H. Kass, E. Klement, T. Waner. Canine monocytic ehrlichiosis: a retrospective study of 100 cases… ExLibris Rapid ILL, Accessed August 7,2022.

[32] Sainz A., Roura X., Miró G., Estrada-Peña A., Kohn B., Harrus S., Solano-Gallego L. Guideline for veterinary practitioners on canine ehrlichiosis and anaplasmosis in europe. https://www.ncbi.nlm.nih.gov/pmc/articles/PMC4324656/, Accessed January 2,2023.

[33] M.A. Sanchez and S.M. Blower. Uncertainty and sensitivity analysis of the basic reproductive rate: tuberculosis as an example. American Journal of Epidemiology, 145(12):1127–1137, 1997.

[34] ScienceDirect. Amblyomma americanum. https://www.sciencedirect.com/topics/medicine-and-dentistry/amblyomma-americanum, Accessed August 7,2022.

[35] Mariah Small and Robert E. Brennan. Detection of rickettsia amblyommatis and ehrlichia chaffeensis in amblyomma americanum inhabiting two urban parks in oklahoma. https://www.ncbi.nlm.nih.gov/pmc/articles/PMC8086404/, 2021. Accessed September 19 2022.

[36] Stanley Coren, Ph.D. How many people don’t walk their dogs? https://www.psychologytoday.com/us/blog/canine-corner/201904/how-many-people-dont-walk-their-dogs, Accessed February 13,2023.

[37] Carlos Alan Torre. Deterministic and stochastic metapopulation models for dengue fever. Arizona State University, 2009.

[38] University of Rhode Island. Lone star tick. https://web.uri.edu/tickencounter/species/lone-star-tick/, Accessed August 7,2022.

[39] P. Van den Driessche and J. Watmough. Reproduction numbers and subthreshold endemic equilibria for compartmental models of disease transmission. Mathematical biosciences, 180(1):29–48, 2002.

[40] Xue Zhang, Bei Sun, and Yijun Lou. Dynamics of a periodic tick-borne disease model with co-feeding and multiple patches. Journal of Mathematical Biology, 82:1–27, 2021.

